# Structural basis of histone H2A lysine 119 deubiquitination by Polycomb Repressive Deubiquitinase BAP1/ASXL1

**DOI:** 10.1101/2023.02.23.529554

**Authors:** Jonathan F. Thomas, Marco Igor Valencia-Sánchez, Simone Tamburri, Susan L. Gloor, Samantha Rustichelli, Victoria Godínez-López, Pablo De Ioannes, Rachel Lee, Stephen Abini-Agbomson, Kristjan Gretarsson, Jonathan M. Burg, Allison R. Hickman, Lu Sun, Saarang Gopinath, Hailey Taylor, Matthew J. Meiners, Marcus A. Cheek, William Rice, Evgeny Nudler, Chao Lu, Michael-Christopher Keogh, Diego Pasini, Karim-Jean Armache

**Author notes:** These authors contributed equally (Jonathan F. Thomas, Marco Igor Valencia-Sánchez).

## Abstract

The maintenance of gene expression patterns during metazoan development is achieved by the actions of Polycomb group (PcG) complexes. An essential modification marking silenced genes is monoubiquitination of histone H2A lysine 119 (H2AK119Ub) deposited by the E3 ubiquitin ligase activity of the non-canonical Polycomb Repressive Complex 1. The Polycomb Repressive Deubiquitinase (PR-DUB) complex cleaves monoubiquitin from histone H2A lysine 119 (H2AK119Ub) to restrict focal H2AK119Ub at Polycomb target sites and to protect active genes from aberrant silencing. BAP1 and ASXL1, subunits that form active PR-DUB, are among the most frequently mutated epigenetic factors in human cancers, underscoring their biological importance. How PR-DUB achieves specificity for H2AK119Ub to regulate Polycomb silencing is unknown, and the mechanisms of most of the mutations in BAP1 and ASXL1 found in cancer have not been established. Here we determine a cryo-EM structure of human BAP1 bound to the ASXL1 DEUBAD domain in complex with a H2AK119Ub nucleosome. Our structural, biochemical, and cellular data reveal the molecular interactions of BAP1 and ASXL1 with histones and DNA that are critical for remodeling the nucleosome and thus establishing specificity for H2AK119Ub. These results further provide a molecular explanation for how >50 mutations in BAP1 and ASXL1 found in cancer can dysregulate H2AK119Ub deubiquitination, providing new insight into understanding cancer etiology.

**One Sentence Summary:** We reveal the molecular mechanism of nucleosomal H2AK119Ub deubiquitination by human BAP1/ASXL1.

## INTRODUCTION

BAP1 is a conserved ubiquitin C-terminal hydrolase responsible for deubiquitinating (DUB) H2AK119Ub, a histone post-translational modification associated with gene-silencing (*1, 2*). BAP1 DUB activity requires the deubiquitinase adaptor domain (DEUBAB) of ASXL1-3 (*3, 4*). In the prevailing model, nuclear BAP1 activates gene expression by deubiquitinating H2AK119Ub and safeguards Polycomb-mediated repression by removing any misplaced H2AK119Ub (*5–8*). Inactivating the catalytic activity of BAP1 abrogates most of its nuclear function, suggesting deubiquitination is central to its role (*5–8*). The *Drosophila* homolog Calypso was identified by genetic screening and characterized as Polycomb Repressive Deubiquitinase Complex (PR-DUB) since it opposes H2A ubiquitination by Polycomb Repressive Complex 1 (PRC1) (*2, 9*). Besides regulating Polycomb repression, PR-DUB activates enhancers and promotes DNA damage repair (DDR) at double-strand breaks (*10–12*). Establishing its importance to human health, BAP1/ASXL1 is frequently implicated in various cancers (*e.g.,* uveal melanoma, malignant mesothelioma) and developmental disorders (*e.g.,* Bohring-Opitz syndrome, myelodysplasia) (*13–18*). Further, mutations in BAP1 are categorized as the “BAP1 tumor predisposition syndrome”, characterized by an increased risk of early onset fast-growing and metastatic cancers (*1, 10*).

Despite its central importance in regulating Polycomb repression, gene activation, and DDR, the primary mechanism for how the PR-DUB specifically deubiquitinates chromatin remains unanswered. While biochemical studies showed that PR-DUB needs nucleosome interactions to be specific for H2AK119Ub and recent crystal structures characterized the architecture and intramolecular interactions in the apo *Drosophila* Calypso/ASX complex, they did not provide a detailed view of how the complex engages chromatin (*3, 19, 20*). Further, these studies have not revealed what determines PR-DUB specificity for H2AK119Ub chromatin, and how cancer-associated mutations in BAP1 and ASXL1 impact its catalytic activity, thereby providing a disease mechanism.

Here, we used a combination of structural, biochemical, and functional approaches to determine the mechanism of nucleosome deubiquitination by human BAP1/ASXL1, one of the most mutated epigenetic complexes among human cancers. We have solved a cryo-EM structure of human BAP1 bound to the ASXL1 DEUBAD domain in complex with a H2AK119Ub nucleosome. BAP1/ASXL1 forms several anchor points on a nucleosome surface and binds H2AK119Ub in a conformation resembling the UCH-L5/RPN13 deubiquitinase (*21*). Specifically, BAP1 interacts with the nucleosome acidic patch (using regions absent in UCH-L5), while ASXL1 interacts with DNA near the nucleosome DNA exit, and both DUB subunits form a DNA clamp that interacts with DNA near the nucleosome dyad. All BAP1/ASXL1 anchor points are mediated through conserved lysine/arginine tracts, which, when mutated, compromise PR-DUB activity *in vitro* and in mouse embryonic stem cells. The position of BAP1/ASXL1 near the dyad interferes with a canonical path of the nucleosomal H2A docking domain, explaining why this portion of histone H2A is disordered in our structure. Based on our results, we propose how the interactions of the DUB complex determine its specificity for H2AK119Ub nucleosomes. Finally, our structural, biochemical, and functional data explain the mechanism of >50 distinct mutations found in cancer, spanning BAP1 and ASXL1 surfaces.

## RESULTS

### Architecture of the BAP1/ASXL1 complex bound to H2AK119Ub nucleosome

To gain insights into the BAP1/ASXL1 deubiquitination mechanism, we reconstituted full-length human BAP1 (residues 1-729) in complex with the DEUBAB domain of ASXL1 (residues 237-390) (Fig. 1A and S1) bound to a nucleosome in which ubiquitin (Ub) residue G76C was conjugated to histone H2AK119C via a non-hydrolyzable dichloroacetate (DCA) linkage (hereafter H2AK119Ub) (*22*). We cross-linked the resulting complex with glutaraldehyde by Gradient Fixation (GraFix) (Fig.S2) (*23*), froze grids, and collected cryo-EM data on a Titan Krios (300 kV) (Table S1). Using these data, we obtained a 3.6 Å resolution cryo-EM map (Fig S3, S4 and Table S2), which we used to unambiguously model the structures of BAP1 (residues 2-60, 69-151, 166-246, 644-713), ASXL1 (residues 248-346), and Ub on the nucleosome (Fig 1B and S5). The cryo-EM map revealed BAP1 bound to the nucleosome in a catalytic conformation, with clearly resolved BAP1-nucleosome, ASXL1-nucleosome, BAP1-Ub, and ASXL1-Ub interfaces (Fig 1B and S5). We used a combination of AlphaFold-Multimer prediction (*24*) and homologous modeling with Drosophila Calypso/ASX crystal structures (PDB ID’s: 6hgc and 6cga) (*19, 20*) to build the model (Fig 1B).

**Fig. 1.**
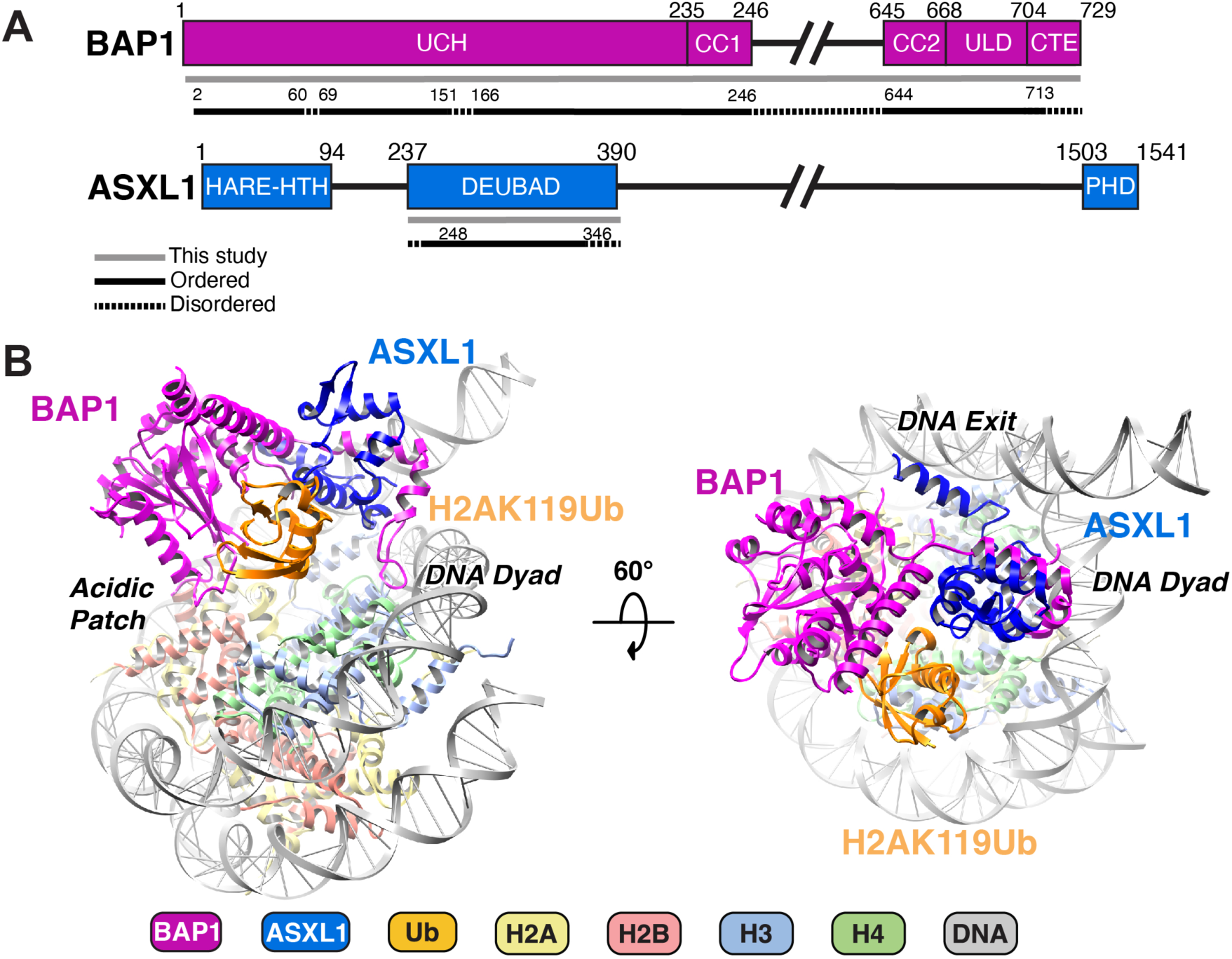
Overview of the structure of BAP1/ASXL1 bound to H2AK119Ub nucleosome. A) Bar diagram representation of BAP1 and ASXL1 domains. Protein sequences included in this study are shown in grey line; those resolved in the structure in black line; and disordered regions in dashed line. B) Two different views of the model for the BAP1/ASXL1-H2AK119Ub nucleosome complex with key anchor points highlighted. The figure is color coded, depicting BAP1 (purple), ASXL1 (dark blue), Ub (Orange), H2A (yellow), H2B (salmon), H3 (light blue), H4 (green), and DNA (grey).

Our structure revealed one molecule each of BAP1 and ASXL1 bound on one side of the nucleosome (Fig 1B and S5). There are four main contact points between BAP1/ASXL1 and the H2AK119Ub nucleosome. BAP1 forms contacts with the acidic patch, DNA and Ub, while ASXL1 interfaces with DNA and Ub (Fig. 1B). These multiple contacts between BAP1/ASXL1 and nucleosome provide a rationale for DUB complex specificity towards H2AK119Ub over other H2A or H2B ubiquitinations (*e.g.* H2AK15ub or H2BK120ub) (Fig. S6).

### The interface of BAP1/ASXL1 with Ub is conserved with other deubiquitinases

Our structure captures ubiquitin in complex with BAP1/ASXL1 on the nucleosome. We used our cryo-EM map and AlphaFold-Multimer prediction to fit the ubiquitin structure (Fig. 2A). We observed that the interfaces between BAP1/ASXL1 and Ub in our model are conserved with UCH-L5/RPN13 (PDB ID 4uem) (Fig. 2A, 2B, S1, and S7) (*21*). In our structure, Ub is sandwiched between the BAP1 UCH and ASXL1 DEUBAD domains (Fig. 2B). BAP1 uses loops Lβ1-β2 and Lα8-β6 to interact with the Ub canonical “I44 patch” (Fig. 2C)(*25*). Our structural analysis and conservation with UCH-L5/RPN13 allowed us to propose the side-chain interactions at the Ub interface (Fig. 2B). BAP1 residues Y33, L35, L230, F228, and I226 are within distance to establish Van der Waals interactions with Ub residues I44, H68, V70, L8, L71, and T9 (Fig. 2C). These residues at the BAP1/ASXL1:Ub interface correspond to those contacting Ub in UCH-L5 and RPN13 (Fig. 2C, 2D, S1, and S7) (*21*), and the ones predicted in Drosophila PR-DUB (*19, 20*). BAP1/ASXL1 also stabilizes Ub by electrostatic interactions on the other face of the “I44 patch”. In this interface, ASXL1 residues R265, H315, and E311, and BAP1 residue E9 are within the distance to establish interactions with Ub residues E24, D39, and R42 (Fig. 2D). This contrasts with other chromatin modifiers that interact primarily with Ub via one of its hydrophobic patches (‘I44 patch’ and ‘I36 patch’)(*25–27*). The BAP1/ASXL1-Ub electrostatic interface is stabilized by potential interactions between ASXL1 ‘NEF motif’ residues H315, F312, and N310, and BAP1 residues R146 (Loop Lα6-α7), R663 and D666 (helix α10) (Fig 2D, S1 and S7). These observations explain why mutations to the NEF motif disrupt catalytic activity by destabilizing Ub binding (Fig S1)(*3, 20, 28*).

**Fig. 2.**
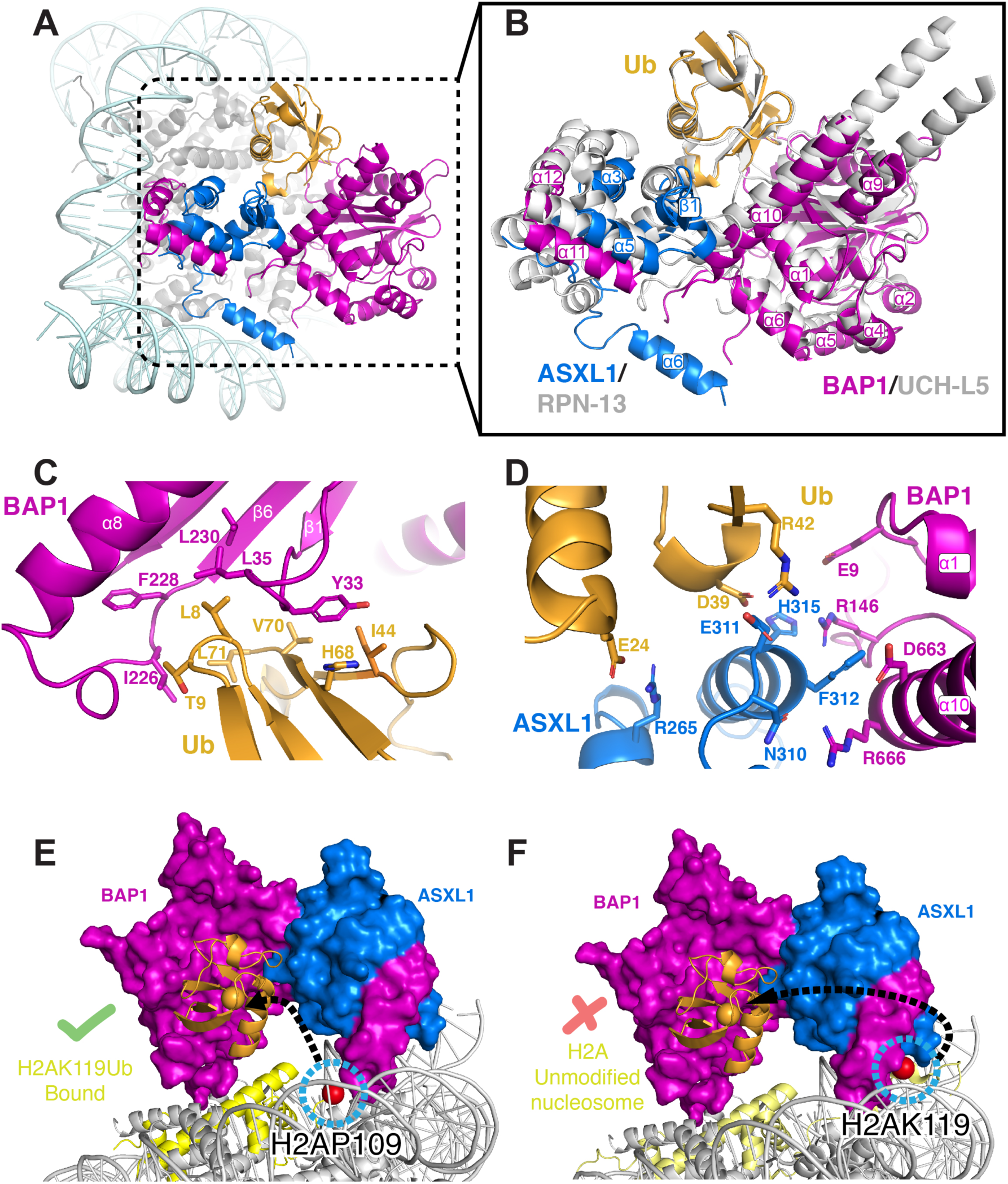
Insights into the mechanism of catalysis by BAP1/ASXL1 and conservation with other deubiquitinases. (A) Position of Ub engaged with BAP1 and ASXL1 on the nucleosome. (B) BAP1/ASXL1/Ub aligned and superimposed with UCH-L5/RPN13/Ub (PDB ID 4uel) (*21*)(grey). (C) Hydrophobic interactions between BAP1 and Ub. (D) Electrostatic interactions between BAP1/ASXL1 and Ub. (E) Space-filling representation of our cryo-EM structure with the last observed residue of H2A (H2AP109; shown as blue circled red sphere). (F) Model with space-filling representation of our Cryo-EM structure superposed with wild type (unmodified) nucleosome structure (PDB ID 1kx5) (*29*) that contains H2A docking domain structured with H2AK119 shown as blue circled red sphere).

### Remodeling of the H2A docking domain allows Ub to reach the catalytic site of BAP1

How does H2AK119Ub reach the catalytic site of BAP1/ASXL1 complex? In our cryo-EM structure, the last ordered residue in the H2A C-terminal tail is P109 (Fig. 2E). The last visible residue of covalently installed Ub, L71, is 37 Å away from P109 and sandwiched below BAP1/ASXL1, which forms an arc-like structure on the nucleosome surface (Fig. 2E). The remainder of the H2A C-terminal tail, up to residue H2AK119 (connected to G76 of Ub via an isopeptide bond and in our structure via a dichloroacetone crosslink between C119 and C76) must be directed to the BAP1 active site for catalysis. This conformation of H2A contrasts with the usual presence of a folded canonical docking domain of H2A in the unoccupied nucleosome (PDB ID 1kx5 and 6wkr) (*29, 30*) bound near the DNA dyad and ordered up to residue K128 (Fig. 2F). This suggests that the H2A docking domain must unfold for H2AK119Ub to reach the catalytic site of BAP1/ASXL1. The distance Ub must relocate defines BAP1 specificity for H2AK119Ub versus H2AK13/15Ub or H2BK120Ub (Fig. S6), since the H2A/H2B histone fold would need to be unwrapped for these to reach the deubiquitinase active site (Fig. S6), revealing why previous studies using a minimal substrate or peptides were unable to explain the specificity for H2AK119Ub on the nucleosome (*3*).

### BAP1/ASXL1 clamps the dyad of the nucleosome

In our structure, BAP1/ASXL1 complex anchors onto the nucleosome surface using three contact points: near the nucleosomal dyad DNA, nucleosomal exit DNA, and the acidic patch. BAP1 residues 672-697 and ASXL1 residues 258-287/311-321 form a helix bundle (Fig. 3A and S1). The helix bundle is formed by ASXL1 helixes αN-3_10_, α1, α2, α3, and α5, folding and wrapping around BAP1 helixes α11 and α12 (Fig. 3A). This helix bundle of BAP1/ASXL1 participates in forming the Ub interface, but also interacts with the nucleosomal DNA near the dyad (Fig. 1B and 3A). The folding of ASXL1 around this part of the ULD region of BAP1 explains previous reports showing that deletion of residues after 670 of BAP1 impairs a complex formation with ASXL1 (*3, 21*). Adjacent to the α12 helix of BAP1, we were able to trace the backbone of BAP1 up until residue 713, where the residues R699, R700, and R701 approach the minor groove near the DNA dyad (Fig. 3B). These three positively charged, well-ordered residues are within the distance to form electrostatic interactions with the DNA phosphates (Fig. S8), and are followed by a C-terminal extension region (CTE) of BAP1 that is needed to recruit PR-DUB to the nucleosomal DNA (*3, 19, 20*). We further explored the roles of R699, R700, and R701 in BAP1 by mutating them to alanines or reversing their charge (Fig. S9). We assembled this mutated BAP1 with ASXL1, and tested catalytic activity of the resulting complex by performing DUB assays on a H2AK119Ub designer nucleosome substrate, observing a significant loss of activity relative to WT (∼84%; Fig. 3A, right, and S9). When we tested nucleosome binding by electromobility shift assay (EMSA), these BAP1 mutations decreased the apparent affinity relative to WT (Fig. 3E, S10 and S11).

**Fig. 3.**
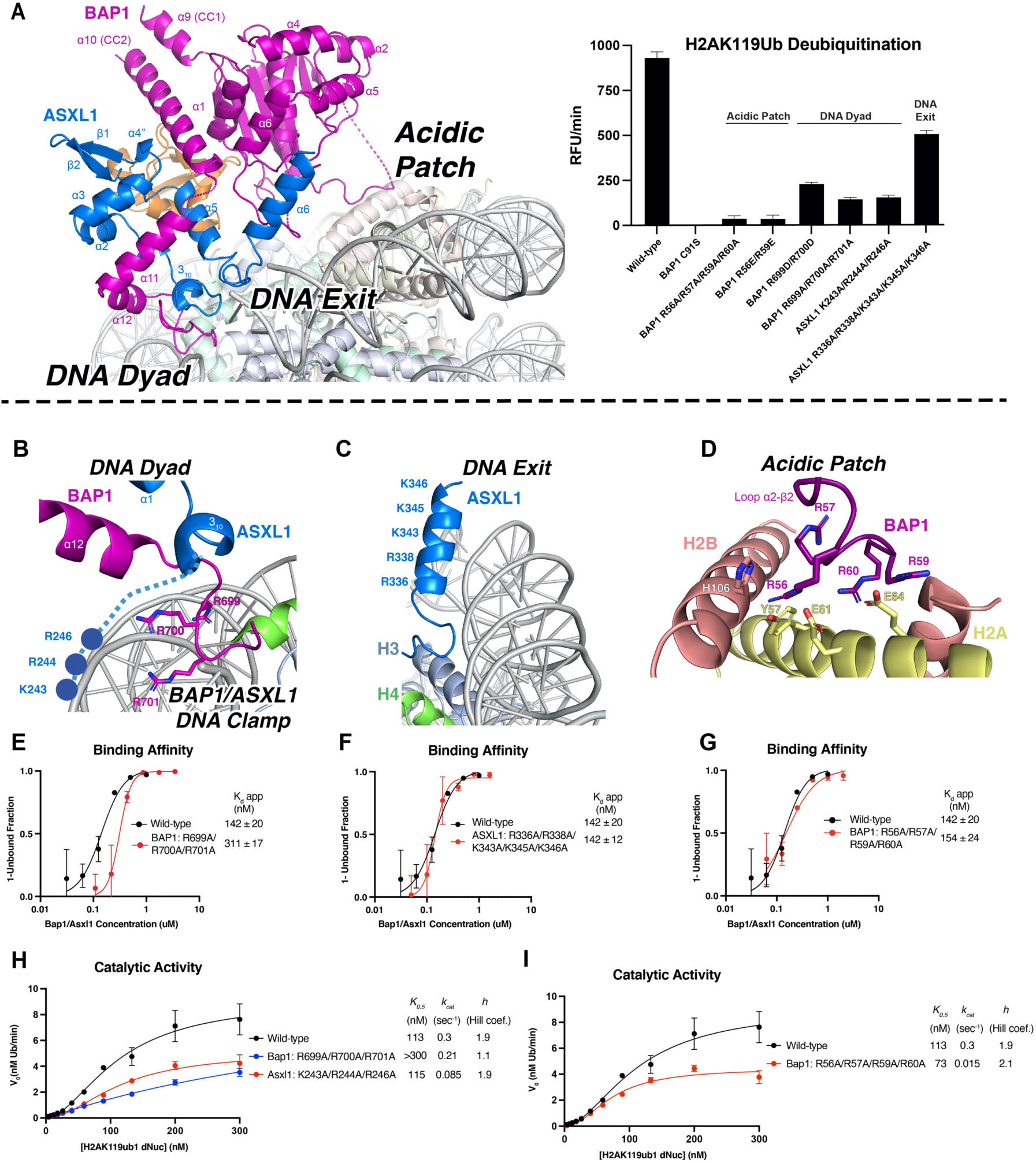
BAP1/ASXL1 interacts with DNA and acidic patch on the nucleosome. (A) Overall architecture of the BAP1/ASXL1-H2AK119Ub nucleosome complex with BAP1/ASXL1 anchor points marked (left). Catalytic activity assays on H2AK119Ub nucleosomes (with a DUB-cleavable native gamma-lysine isopeptide linkage) and wild type BAP1/ASXL1, the catalytic inactive mutant of BAP1 (C91S) /ASXL1, and mutations to the nucleosomal anchors (right). (B) Close up view of BAP1/ASXL1 DNA clamp contacting the nucleosome near the DNA dyad. (C) Close up view of the ASXL1 DEUBAD α6 helix, projecting a stretch of basic residues towards a nucleosome DNA exit. (D) Close up view of the BAP1 R-finger interacting with the acidic patch. Quantified results from electromobility shift assay (EMSA) of wild type and mutants of BAP1 DNA clamp (E), ASXL1 DNA exit (F), and BAP1 interaction with the acidic patch (G). Each data point and error bar indicate the mean ± SD from three independent experiments. The standard errors of dissociation constants (Kd) are indicated. Michaelis-Menten curves of BAP1/ASXL1 DNA clamp mutations (H) and the acidic patch mutations (I) on H2AK119Ub nucleosomes. Please see Supplementary Text for extended information on figures 3A, H and I.

Immediately before the αN-3_10_ helix, the ASXL1 N-terminal tail also contains a patch of basic residues (R243, R244, and K246) that we hypothesized can form a clamp together with the basic residues of BAP1 near the helix bundle (Fig. 3A, 3B and S1). We mutated these three residues in ASXL1 to alanines, assembled a complex with BAP1, and again observed a significant loss of DUB activity (∼83%) to H2AK119Ub nucleosomes (Fig. 3A, right). We termed this region the ‘DNA clamp’ since the basic residues of BAP1 (R699, R700, and R701) and ASXL1 (R243, R244, and K246) provide extensive interactions with the double helix near the DNA dyad (Fig. 3B). To further explore enzymatic mechanism, we performed kinetic assays using wild-type BAP1/ASXL1 and DNA clamp mutants (BAP1 R699A, R700A, and R701A; ASXL1 R243A, R244A, and K246A). The BAP1 mutant complex compromised catalytic activity where *K*_0.5_ was increased (>300 nM vs 113 nM WT) and *k*_cat_ was slightly reduced (0.21 vs 0.3 sec^−1^ WT) (Fig. 3H). In contrast, the ASXL1 mutant complex showed no effect on binding affinity but reduced *k*_cat_ (0.085 sec^−1^ vs 0.3 sec^−1^ WT) (Fig. 3H, S10, S11). The observation that mutations of BAP1 or ASXL1 residues in the DNA clamp have a significant impact on catalytic activity supports the clamp’s importance in the deubiquitination mechanism of H2AK119Ub (Fig. 3A).

### ASXL1 interacts with the DNA at the exit of the nucleosome

Our cryo-EM map showed an extra density emerging from the helix bundle and DNA clamp and approaching the DNA exit. During the cryo-EM data processing, this point of contact was only seen when BAP1/ASXL1 was in the catalytic conformation and Ub was stabilized (Fig. S4). The rigid body fit of this region allowed us to assign it to the ASXL1 helix α6 (Fig. 3C), corresponding to Drosophila ASX α6 (PDB ID 6hgc)(*19*). Even though the map in this region does not allow to build side-chain contacts, this region of ASXL1 contains a stretch of basic residues (R336, R338, K343, K345, and K346), which we hypothesized may interact with the DNA backbone (Fig. 3C). We mutated these basic residues in ASXL1 (R336, R338, K343, K345, and K346) to alanines, assembled with WT BAP1, and tested DUB activity of the resulting complex. Relative to WT, we observed a moderate reduction of catalytic activity (∼46%) (Fig. 3A, right), with no change in the binding affinity (Fig. 3F), supporting the importance of this interface for nucleosome interaction.

### BAP1 binds the acidic patch with an arginine finger

Another stable anchor between BAP1/ASXL1 and the nucleosome in our cryo-EM map is with the acidic patch, judging by a better resolution of this interface (Fig. S8, S12, and S13). Here, we were able to assign the side chains of the BAP1 loop Lβ2-α2 residues (R56, R57, R59, and R60) (Fig. 3D, S1 and S8), which formed electrostatic interactions with the acidic patch residues E61, E64, and Y57 of H2A and H106 of H2B (Fig. 3D). This R-finger is the classical interface between chromatin modifiers and the nucleosomal H2A-H2B dimer (*26, 31–36*). These four arginines are conserved between mammalian BAP1 and *Drosophila* Calypso, but not in UCH-L5, another DUB which does not deubiquitinate a nucleosome substrate (Fig. S1). We tested the functional significance of these arginines (R56, R57, R59, and R60) by mutating them to alanines or glutamic acid, complexed with ASXL1, and performing DUB or EMSA assays with H2AK119Ub nucleosomes. Relative to WT, these BAP1/ASXL1 mutants showed almost a complete loss (∼96%) of catalytic activity (Fig. 3A, right) but no change to binding affinity (154 nM vs 142 nM) (Fig. 3G). Determination of the kinetic constants of the BAP1 loop Lβ2-α2 mutant complex revealed a significant reduction in turnover number (*k*_cat_= 0.015 vs 0.3 sec^−1^) and a slight change of the *K*_0.5_ (Fig. 3I). These results show that binding affinity is not perturbed when disrupting the interaction with the acidic patch, in agreement with a previous report (*3*). However, we demonstrated that this interface is critical for PR-DUB catalytic activity, closely mirroring the role of arginine finger-acidic patch interactions in the activity of other chromatin enzymes (*26, 31, 32, 37*).

In conclusion, all anchor points contributed to H2AK119-directed deubiquitination activity, and mutating each interface disrupted catalysis (Fig. 3A, right). Our cryo-EM structure suggests a mechanistic model where BAP1/ASXL1 deubiquitinates chromatin by anchoring to the nucleosomal acidic patch and DNA at the dyad and exit. Complementary *in vitro* assays (binding affinity, catalytic activity, and kinetics) indicate that these anchor points contribute to enzyme:substrate engagement and catalysis with varying degrees of importance.

### BAP1/ASXL1-nucleosome interactions are required for H2AK119Ub deubiquitination in mESCs

We next sought to examine the *in vivo* functional importance of the molecular contacts identified in our structural and biochemical analyses. To determine whether the residues identified by our analyses directly regulate BAP1 H2AK119Ub DUB activity, we stably re-expressed BAP1 R699E/R700E (DNA dyad), BAP1 R56E/R59E, BAP1 R56A/R57A/R59A/R60A (acidic patch) in BAP1 KO ESC in parallel to either a wild-type or BAP1 catalytic inactive form (C91S). All BAP1 mutants expressed at comparable levels with respect to the BAP1 WT counterpart (Fig. 4A). As we previously reported, the re-expression of WT BAP1 in KO ESC efficiently restored physiological H2AK119Ub levels (*6*). However, consistent with our *in vitro* data, the expression of all distinct BAP1 mutations failed to restore physiological H2AK119Ub levels similar to a catalytically inactive form of BAP1 (Fig. 4A). Together, these results confirm that the critical residues involved in BAP1/ASXL1 H2AK119Ub nucleosome recognition (Fig. 4B) are required for efficient deubiquitination *in vivo*.

**Fig. 4.**
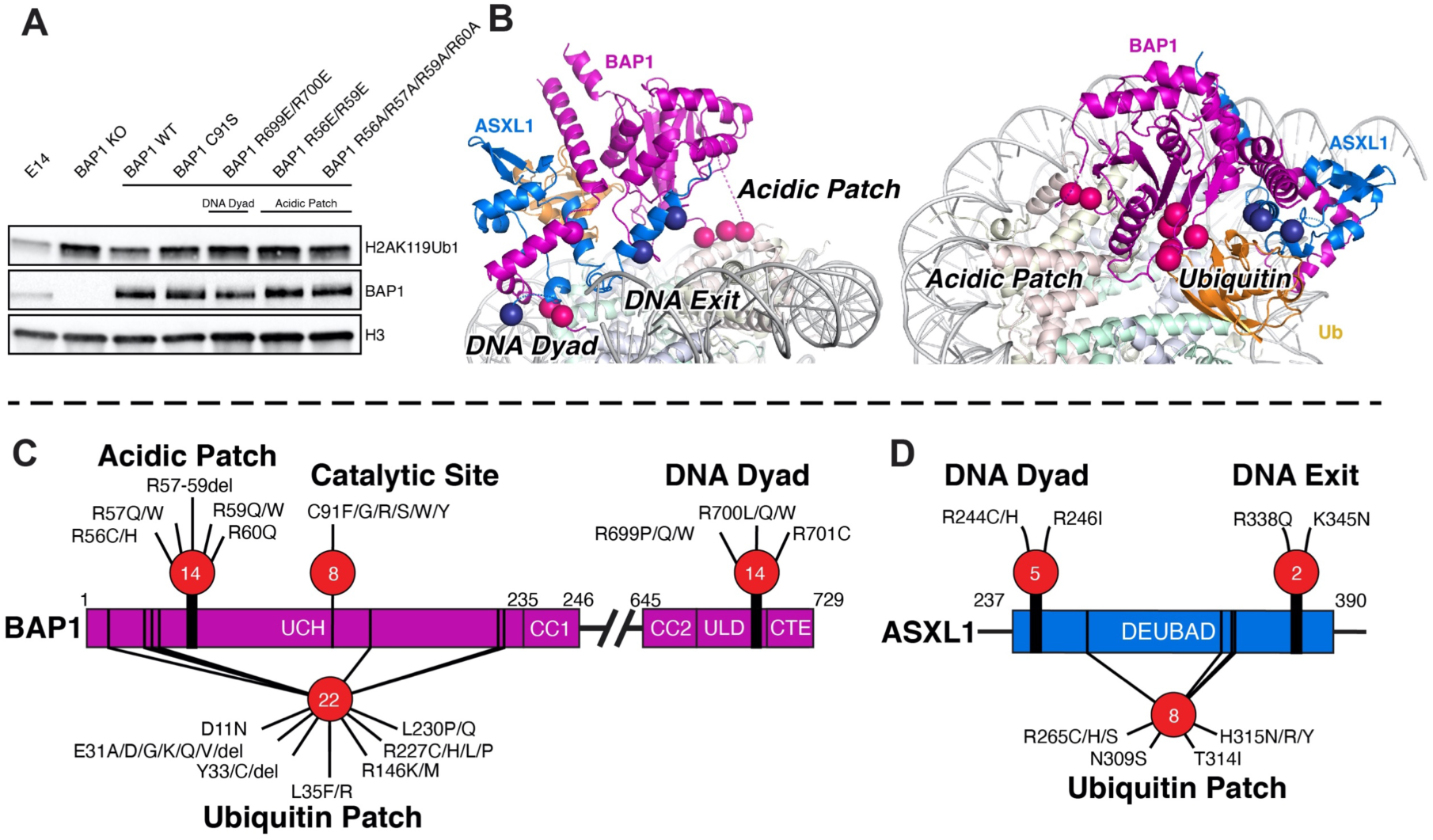
BAP1/ASXL1-nucleosome contacts are extensively mutated in cancers. (A) Western blot analysis using the indicated antibodies of total protein extracts from BAP1 WT (E14) or KO mESC with stable expression of WT and various BAP1 mutations (C91S, R699E/R700E; R56E/R59E; R56A/R57A/R59A/R60A). (B) Structure of the BAP1/ASXL1-H2AK119Ub nucleosome complex highlighting three key nucleosome anchors (left) and Ub interactions (right) with cancer mutations shown as colored spheres. Bar diagram showing cancer-associated point mutations and deletions to BAP1 (C) and ASXL1 (D) at nucleosomal anchors and the Ub patch shown in (B) identified in several cancers that can be mechanistically explained by studies presented here. The number of unique cancer types at each interface is shown inside the red spheres.

### Cancer mutations decorate BAP1/ASXL1-nucleosome interfaces

BAP1/ASXL1 is one of the most frequently mutated chromatin-modifying enzymes in cancers, with its disease relevance further highlighted by BAP1 mutations being classified as “BAP1 Predisposition Tumour Syndrome” (*38*). BAP1 and ASXL1 are decorated with mutations across the entire polypeptide chain, with most driver mutations being truncations and more than 99% of variants of uncertain significance (VUS) being missense mutations. Most truncation mutations disrupting function are easily explained by loss of domains or nuclear localization signals (*1*). In contrast, gain of function truncations that decorate the middle of ASXL1 were shown to stabilize BAP1 on chromatin and reduce BAP1 degradation (*39, 40*). However, the abundant VUS missense mutations often remain mechanistically unexplained – especially when a protein structure is unavailable (S14 and S15). The Drosophila PR-DUB crystal structures gave context for cancer-associated intramolecular mutations between BAP1-ASXL1-UB, but left the interactions of the complex with nucleosomes unexplained (*19, 20*). Our structure provides context for VUS missense mutations that define deubiquitination activity on nucleosomes (Table S3 and S4). We searched cBioPortal (https://www.cbioportal.org) and used a curated set of non-redundant studies to both tabulate a list of all cancer mutations across BAP1 and ASXL1 (Table S3 and S4) and annotate the subset of frequent mutations at the Ub interface and the DNA dyad, DNA exit, and acidic patch interfaces (Fig. 4B). On BAP1, deletion or non-conservative mutations of the acidic patch–interacting residues R56–60 or the DNA dyad interface residues R699–701 are each found in 14 different cancers (Fig. 4C). For ASXL1, mutations of the DNA dyad residues R244 and R246 are found in five different cancers (Fig. 4D), while the DNA exit residues R338 and K345 are found in colorectal adenocarcinoma and cholangiocarcinoma (Fig. 4D). Taken together, we explain 1% of ASXL1 and 5% of BAP1 VUS missense mutations at key nucleosome interfaces, providing key insights into the potential deregulation of BAP1/ASXL1 in 35 cancers ranging from adenoid cystic carcinoma to uveal melanoma. cBioPortal functional prediction and our structural, biochemical, and cellular studies indicate all these mutations lead to a loss of function where BAP1/ASXL1 fails to effectively deubiquitinate H2AK119Ub, and thus causes aberrant accumulation of this chromatin modification.

## Discussion

Our data support a model where BAP1/ASXL1 deubiquitinates H2AK119Ub chromatin by anchoring to the nucleosomal acidic patch, DNA dyad, and the DNA exit regions using conserved arginine and lysine tracts. Fitting the prevailing model for chromatin-modifying proteins(*32, 41*), the ability of BAP1 to fine-tune deubiquitination activity requires interactions with the nucleosome acidic patch. Previous reports showed that ASXL1 increases the affinity of PR-DUB for ubiquitin, and revealed the role of BAP1 CTE region in enabling PR-DUB binding to nucleosomal DNA (*3, 19*). Here we show the molecular interactions of BAP1/ASXL1 that direct the PR-DUB complex to the nucleosome and enzymatically characterize these interfaces using a defined H2AK119Ub nucleosome substrate. We describe a novel functional element, the BAP1/ASXL1 DNA clamp containing a region of the helix bundle of ASXL1 folded around BAP1 and connecting to the CTE. We show that BAP1 residues that form the clamp are critical for efficient deubiquitination of nucleosome substrate by directing the PR-DUB to its catalytic position on the DNA dyad and clarify the importance of ASXL1 for DUB activity (*3*). Our cryo-EM structure also provides a rationale as to how PR-DUB mediates substrate specificity for nucleosomal H2AK119Ub over H2AK13/K15Ub and H2BK120Ub - in the latter cases the ubiquitylated residue cannot reach the BAP1/ASXL1 active site without extensive remodeling of the H2A/H2B dimer (Fig. 2E, 2F and S6). There is a functional redundancy in deubiquitination activity between ASXL1, 2, and 3 (*3*). Our structure reveals that the residues in ASXL1 involved in nucleosome binding are conserved in ASXL2 and 3, explaining their functional redundancy in the PR-DUB (Fig. S1). Lastly, our structure explains VUS cancer mutations that may disrupt deubiquitination activity by abrogating the interaction between BAP1/ASXL1 and substrate nucleosomes.

We showed the residues R699–R701 of the BAP1 DNA clamp to form important interactions with the DNA dyad and lead to the CTE, which extends towards the DNA exit. The deletion of residues of the CTE of BAP1 after H710 has been reported to reduce the recruitment of PR-DUB to endogenous nucleosomes (*3, 19, 20*). We hypothesize that the non-specific interactions of the positively charged CTE with DNA have two important roles. 1) They participate in the proper folding/positioning of the DNA clamp near the DNA dyad to allow PR-DUB to adopt an active conformation. 2) In a similar fashion as the disordered basic residues of the N-terminal portion of ASXL1, the CTE of BAP1 would help recruit the BAP1/ASXL1 clamp to the DNA. The strong recruitment to nucleosomal DNA via a tract of basic residues and establishing non-specific interactions may be a common feature of chromatin modifiers (*27, 31, 37, 42*).

In this study, we primarily discuss a cryo-EM map characterized by a stable binding of the three anchors of BAP1/ASXL1 with the nucleosome acidic patch, DNA clamp, and DNA exit regions. The particles used to calculate this map also contain the Ub stabilized in its catalytic conformation (Fig. S4 and S12). This map showed ‘the best’ defined interfaces, which we interpret as the most stable conformation of BAP1/ASXL1 on the H2AK119Ub nucleosome. In addition, during data processing, we observed 3D classes with alternative conformations of BAP1/ASXL1 on nucleosomes (Fig. S4 and S12). In the 3D classification analysis, some of the classes presented higher flexibility in the domains of BAP1/ASXL1 (as evidenced by lower resolution maps) and did not show ubiquitin stabilization in its catalytic conformation (Fig. S4 and S12). 35% of the particles showed the BAP1/ASXL1 DNA clamp/CTE binding to the DNA exit instead of the dyad (alternative conformation). Based on our 3D classification analyses, we propose that when the PR-DUB is recruited to chromatin, it samples these alternative conformations to find the catalytic position. The catalytic position is established by pivoting on the nucleosome acidic patch to position the clamp on the dyad.

The BAP1/ASXL1 DNA clamp on the dyad, H2A docking domain in the canonical nucleosome conformation, and histone H1.4 all occupy almost the same location (*43, 44*). The linker histone H1.4 binds asymmetrically to the linker DNA depending on the nucleosome stacking in the chromatin (PDB ID 7pfv)(*43*). Thus, the BAP1/ASXL1 DNA clamp may clash with H1.4 bound at the same side of the DNA linker (Fig. S12). We hypothesize that H1.4 binding in this mode would force BAP1/ASXL1 to engage in the alternative conformation at the DNA exit instead of the DNA dyad. In the corollary, this could explain the H1 enrichment on chromatin after BAP1/ASXL1 deletion or H2AK119 deubiquitination (*6, 45*).

Similar to other chromatin complexes, such as Dot1 or COMPASS, the observed conformational flexibility of BAP1/ASXL1 on the nucleosome could be regulated. Both Dot1 and COMPASS are stabilized in the catalytic conformation on the nucleosome by interactions with histone modifications such as histone ubiquitination or acetylation (*26, 27*). In the case of BAP1/ASXL1, histone modifications, histone variants, linker histones, and co-factors could play stabilizing / regulatory roles and should be the focus of subsequent studies.

## Acknowledgements

We thank colleagues for the generous supply of materials (see Methods); Dr. Bing Wang for helping with data collection at NYU cryo-EM Shared Resource. We thank Dr. Cohen and Dr. Yao’s labs for proving the tUI sensor. We thank Michael Costantino, Kevin Yie, and staff of the HPC Core at NYU Langone Health for computer access and support; and Dr. Peter Hare for critical reading of the manuscript. We thank all members of the Armache laboratory for helpful suggestions and discussions.

## Funding

Work in the Armache laboratory is supported by grants from the Mark Foundation for Cancer Research, and the National Institutes of Health R01GM115882 and R01CA266978. Work in the Pasini laboratory was supported by the World Wide Cancer Research (22–0027); the Italian Association for Cancer Research, AIRC (IG-2017-20290 and IG 2022-27694); and by the European Research Council, ERC (EC-H2020-ERC-CoG-DissectPcG: 725268). S.T work is supported by grants from AIRC (MFAG 2021-26131) and Fondazione Cariplo (Cariplo Giovani 2020-3576). S.R. is a PhD student within the European School of Molecular Medicine (SEMM). Work in the Lu laboratory is supported by R35GM138181. Work in *EpiCypher* is supported by NIH grant R44 GM119893.

## Author Contributions

J.T, M.I.V.-S, and K.-J.A. conceptualized and designed the study. J.T., M.I.V.-S., V.G.L., P.D., R.L. and S.A.A. performed the structural and biochemical experiments. S.L.G., J.M.B., A.R.H., L.S., S.G., H.T., M.J.M., M.A.C. and M-C.K. made designer nucleosomes and performed activity and kinetic assays. S.T., S.R., and D.P. performed studies in mESCs. W.R. contributed to data collection and processing. All the authors above contributed to data analysis, interpretation, and writing of the manuscript with input from K.G., E.N., and C.L.

## Competing interests

*EpiCypher* is a commercial developer and supplier of reagents (*e.g.* PTM-defined semi-synthetic nucleosomes; dNucs) used in this study.

## Data and materials availability

All data are available in the manuscript or the supplementary materials. The structure models and the cryo-EM density maps have been deposited in PDB and EMDB with accession codes: XXXX, EMD-XXXXX. Further inquiries and requests for resources and reagents should be directed to and will be fulfilled by the Lead Contact, Karim-Jean Armache.

## Materials and Methods

### Cloning and mutation of BAP1 and ASXL1

Full-length human BAP1 and the deubiquitinase adaptor domain (DEUBAD) of ASXL1 (amino acids 237-390) were cloned into a pFastBac Dual vector by GenScript. ASXL1 was subcloned into pET24a vector for *E. coli* expression. BAP1 was modified to express alone in SF9 from the pFastBac Dual vector. The mutants described in this study were generated using NEB Q5 Site-Directed Mutagenesis Kit.

### Protein expression and purification of BAP1 and ASXL1

BAP1/ASXL1 and BAP1 were expressed in SF9 insect cells (clonal isolate from PLB-Sf-21-AE1) following the Bac-To-Bac Baculovirus Expression System established protocol (Invitrogen). ASXL1 was expressed in *E. coli* RIL by IPTG induction following established protocols (*19*).

BAP1/ASXL1 complex and BAP1 alone were purified as previously with adaptations (*28*). First, cells were suspended in Resuspension Buffer A (20 mM HEPES pH 8, 150 mM NaCl, 5% glycerol, 0.5 mM TCEP, Roche EDTA free protease inhibitor) and lysed using an emulsifier (AvestinEmulsiflexC3). Lysate was precleared by spinning at 7,500 RCF for 20 minutes. BAP1 was purified by Strep-Tactin XT affinity chromatography resin (Iba). The lysate was incubated with Strep-Tactin XT resin, washed with Resuspension Buffer A, and sequentially washed with Buffer A supplemented with 1M NaCl and 2M NaCl. BAP1 was cleaved from resin by incubating overnight with 600 µg of 3C protease. Elution fractions were dialyzed overnight in Buffer A (20 mM HEPES pH 8, 150 mM NaCl, 5% glycerol, 0.5 mM TCEP), then passed through a HiTrap Q HP 5ml column, and eluted by an increasing gradient of Buffer B (20 mM HEPES pH 8, 1 M NaCl, 5% glycerol, 0.5 mM TCEP). Lastly, BAP1 or BAP1/ASXL1 were further purified on gel filtration column (HiLoad 16/60 Superdex 200 or Superose 6 Increase 10/300 GL (Cytiva)) using gel filtration Buffer C (10 mM HEPES pH 7.5, 150 mM NaCl, 10% Glycerol, 0.5 mM TCEP). BAP1 or BAP1/ASXL1 fractions were pooled, concentrated, and snap-frozen.

ASXL1 was purified as previously (*19*) with some modifications. ASXL1 was expressed in *E. coli* BL21 (DE3) and collected by centrifugation at 7,500 RCF for 15 minutes and then suspended in Lysis Buffer D (20 mM HEPES pH 8, 200 mM NaCl, 5 mM imidazole, 0.5 mM Imidazole, 1x Roche EDTA free protease inhibitor). Cells were lysed using an emulsifier (AvestinEmulsiflexC3) and the lysate was cleared by spinning at 7,500 RCF for 20 minutes. ASXL1 was purified by using Ni-NTA resin (Qiagen). The resin was washed with Lysis Buffer D, and sequentially washed with buffer D supplemented with 1 M NaCl, and returned to Buffer D. ASXL1 was eluted by cleaving from resin with 3C protease. Elution fractions were concentrated using a 10K MW cutoff Centricon concentrator, and passed over a HiLoad Superdex 75 16/60 pg (Cytiva) using Gel Filtration Buffer C (10 mM HEPES pH 7.5, 150 mM NaCl, 10% Glycerol, 0.5 mM TCEP). ASXL1 containing fractions were pooled, concentrated, and snap frozen.

### BAP1/ASXL1 complex assembly

Purified BAP1 was co-incubated with an excess of ASXL1 for 1 hour at 4°C before injecting into the Superose 6 Increase 10/300 GL (Cytiva) using Gel Filtration Buffer C (10 mM HEPES pH 7.5, 150 mM NaCl, 10% Glycerol, 0.5 mM TCEP). Peak fractions containing BAP1/ASXL1 were pooled, concentrated, and snap frozen.

### Expression and purification of Ubiquitin

pET-His-Ub G76C was a generous gift from Dr. Tingting Yao (*22*). Ubiquitin plasmid DNA was transformed into *E. coli* SoluBL21™ (Amsbio, Cat# C700200) competent cells and grown in 2xYT-Amp Media. Ub G76C (Ub G76C) was expressed as soluble protein by inducing with 0.4 mM IPTG for 4 hours at 37°C upon the culture reaching OD600 = 0.4-0.6. Bacteria cells were harvested and lysed (AvestinEmulsiflexC3). Protein was purified through Ni-NTA agarose beads (Qiagen) (Lysis Buffer: 300 mM NaCl, 50 mM Tris pH 8.0, 10 mM Imidazole, 5 mM BME, 1x Protease Inhibitor / Elution Buffer: 300 mM NaCl, 50 mM Tris pH 8.0, 300 mM Imidazole, 5 mM BME) followed by HiTrap Q HP (GE Healthcare) liquid chromatography column (Buffer A: 50 mM NaCl, 20 mM Tris pH 8.0, 0.2 mM EDTA, 10 mM BME / Buffer B: 1 M NaCl, 20 mM Tris pH 8.0, 0.2 mM EDTA, 10 mM BME). Purified Ub was then dialyzed against water supplemented with 1 mM acetic acid followed by flash freezing in liquid nitrogen and lyophilized using a Sentry lyophilizer (VirTis).

### Purification of Widom 601 DNA 187 bp

A plasmid containing the Widom 601 nucleosome positioning sequence extended to 187 bp and flanked by EcoRV restriction sites (*36*) was transformed into *E. coli* DH5α (ThermoFisher) and grown in 2xYT-ampicillin media overnight at 37°C. The 601 DNA fragment was excised using EcoRV and purified as previously (*46*).

### Expression and purification of wild-type *Xenopus* histones and H2AK119C

Plasmids containing wild-type *Xenopus* histones were a generous gift from Dr. Karolin Luger, and mutant histone H2AK119C was generated using the Q5 mutagenesis kit (NEB). Briefly, each histone was expressed in *E. coli* Rosetta (DE3) cells (Novagen), extracted from inclusion bodies, and purified sequentially by size exclusion and anion chromatography as previously (*46*). Purified histones were freeze dried using a Sentry lyophilizer (VirTis).

### Ubiquitination of histone H2AK119C (non-labile for cryo-EM)

We followed a previously published protocol (*22*). Briefly, lyophilized Ub G76C and histone H2AK119C were re-suspended (Re-suspension buffer: 10 mM Acetic acid, 7M Urea-Deionized) and mixed in the ratio of 2:1. Sodium Tetraborate, Urea, and TCEP were added to achieve final concentrations of 50 mM, 6M and 5 mM, respectively. The mixture was incubated at room-temperature for 30 minutes. Then, an amount of crosslinker (1,3-dichloroacetone (Sigma) diluted in N,N’-dimethyl-formamide (Sigma)) equal to one-half molar ratio of total sulfhydryl groups was added to the solution and incubated on ice for additional 30 minutes. The reaction was stopped by addition of BME to a final concentration of 5 mM. Then, the solution was diluted 10 times with denaturing binding buffer (Denaturing Binding Buffer: 50 mM Sodium phosphate (NaPi), 50 mM Tris pH 8.0, 300 mM NaCl, 6 M Urea, 10 mM Imidazole, 5 mM BME) and purified through Ni-NTA agarose beads (Qiagen) (Denaturing Elution Buffer: 50 mM NaPi, 50 mM Tris pH 8.0, 300 mM NaCl, 6 M Urea, 250 mM Imidazole, 5 mM BME). Purified Ubiquitinated-H2AK119C (H2AK119Ub) was dialyzed into water supplemented with 1 mM BME and lyophilized using Sentry lyophilizer (VirTis).

### Reconstitution of nucleosomes

Nucleosome substrates were assembled as described(*26, 31*). Equimolar amounts of each lyophilized 4 histones were dissolved in unfolding buffer (6 M guanidinium hydrochloride, 20 mM Tris, pH 7.5, 5 mM DTT), mixed and dialyzed into refolding buffer. Octamers were purified through size exclusion chromatography Superdex 200 column (GE healthcare) in refolding buffer (10 mM Tris, pH 7.5, 2 M NaCl, 1 mM EDTA, 5 mM BME). Nucleosomes were assembled by combining equimolar ratios of purified Widom 601 DNA and histone octamers, and dialyzing the mix overnight with gradient salt dialysis using a peristaltic Rapid Pump (Gilson). Assembled nucleosomes were purified through a Resource Q ion-exchange column (GE Healthcare). Finally, purified nucleosomes were dialyzed into TCS buffer (20 mM Tris-HCl pH 7.5, 1 mM EDTA, 1 mM DTT), concentrated, and stored at 4°C until use.

### Designer Nucleosomes (dNucs)

PTM-defined nucleosomes (all from *EpiCypher*: rNuc (unmodified) <u>#16-0006</u>; H2AK15ub1 <u>#16-0399</u>; H2AK119ub1, <u>#16-0395</u>; H2BK120ub1, <u>#16-0396</u>) were assembled through salt-dialysis of semi-synthetic histones with 5’ biotinylated DNA (147bp of 601 nucleosome positioning sequence) as previously (*47, 48*) and confirmed by SDS-PAGE, immunoblotting, and mass spectrometry (as appropriate). All Kub1 histones contain a native gamma-lysine isopeptide linkage cleavable by deubiquitinating (DUB) enzymes (*49*).

### Free Ub system of detection

tUI free Ub sensor (*50*) was generously provided by Robert Cohen and Tingting Yao (Colorado State University). Briefly, the sensor is a fusion protein containing ubiquitin binding domains from ubiquilin-1 (UQ1; aa541-586) and isotranspeptidase T (IsoT; aa163-91) and relies on the fluorescence change of a single covalently attached Atto-532 when it engages free ubiquitin (as released by DUB activity on a Kub1 dNuc).

### Gradient fixation (GraFix) of BAP1/ASXL1-H2AK119Ub nucleosome

H2AK119Ub nucleosome on 187bp DNA was saturated with BAP1/ASXL1 and dialyzed in 5% Glycerol Buffer A2 (20 mM HEPES pH 7.5, 50 mM KCl, 1 mM DTT, 5% glycerol). BAP1/ASXL1-nucleosome complex was crosslinked using Gradient Fixation (GraFix) as previously (*23*). Gradients were produced (Gradient Master, Biocomp instrument) using a gradient range of 10% glycerol buffer B2 (20 mM HEPES pH 7.5, 50 mM KCl, 1 mM DTT, 10% glycerol) to 40% glycerol buffer C2 (20 mM HEPES pH 7.5, 50 mM KCl, 1 mM DTT, 40% glycerol, 0.1% glutaraldehyde). Gradients were spun at 30,000 RPM for 16 hours at 4°C, fractionated, and remnant glutaraldehyde quenched with 100 mM Tris-Cl pH 7.5 (final). Fractions were analyzed by 4.5% native PAGE (0.2X TBE) and those of interest pooled and dialyzed in cryo-EM Buffer (20 mM Tris pH 7.5, 50 mM KCl, 1 mM DTT, 1% glycerol). Final samples were concentrated and used for grid preparation.

### Grid preparation for cryo-EM

Cryo-EM grids of the BAP1/ASXL1-H2AK119Ub nucleosome complex were prepared with the following established protocol (*51*). Briefly, Quantifoil R 1.2 µm hole size Au 300 mesh grids were glow discharged for 25s at 15 mA using a PELCO easiGlow. 3 µl of the sample at 0.1 mg/mL concentration were applied to grids using a FEI Vitrobot Mark IV (FEI Company), at 4°C and 100% humidity. Grids were blotted using filter paper (55/20 mm diameter, TED PELLA) for three seconds with a force of 3 and plunge-frozen in liquid ethane cooled to liquid nitrogen temperatures.

### Cryo-EM data processing

Data was collected using a Titan Krios G3i using Leginon software (*52*). All images were recorded using a K3 Summit Gatan direct electron detector camera at a nominal magnification of 105,000X, calibrated physical pixel size of 0.825 Å/pixel (0.4215 in super-resolution Å/pixel) using an energy filter width of 20. The images were collected with an exposure of 0.04 seconds per frame; the total exposure time was 2 seconds for the BAP1/ASXL1-H2AK119Ub complex, for a total of 50 frames. Per-frame exposure rate was 1.1424 e/Å^2^ leading to a total accumulated electron exposure of 57.12 electrons per Å^2^ on the specimen. All the sample images were recorded with a calibrated defocus in the range from −1.2 to −2.4 μm. Movie stacks, acquired in super-resolution mode, were corrected for global and local motions (in 11×9 patches) using UCSF MotionCor2 v1.2.1 (*53*) ran through Relion context (*54*), resulting in the dose-weighted sums. We imported the motion-corrected dose-weighted images to CryoSPARC (*55*) and performed Patch CTF Estimation. We used ‘blob-based’ automated particle picking to obtain 11,876,334 particles, and selected 5,510,142 particles for subsequent processing. Particles were then extracted and subjected to reference-free 2D classification into 50 classes in CryoSPARC. A small number of particles (∼265K) were selected from the seven best-looking reference-free 2D classes. We then used Cryosparc’s “Ab initio” option to generate a 3D initial model from the selected small dataset. This yielded a reconstruction with a clearly defined nucleosome density, which was then used as a template in 3D refinement. For data “in silico” homogenization of our data, we employed CryoSPARC’s “Ab initio” and “Heterogeneous refinement”. After 3D refinement, we used focused classification to improve the density of BAP1/ASXLl in the maps, the data was subject to focused classifications. In the BAP1/ASXL1-H2AK119Ub complex, we centered a spherical mask with a 60 Å pixels radius on the BAP1/ASXL1 moiety, which resulted in a clearly resolved BAP1/ASXL1 on the nucleosome at 5.06 Å overall resolution. These particles were subject to a variability analysis, using a similar mask centered on the BAP1/ASXL1 moiety, using Principal Component Analysis with 5 modes and an output of 10 clusters, finding one cluster that contained a better-resolved BAP1/ASXL1 clamp-DNA and BAP1/ASXL1-Ub interfaces at 4.35 Å after extraction of the particles. Using the best-scoring particles, we ran Non-Uniform Refinement, which yielded the BAP1/ASXL1-H2AK119Ub complex at 3.6 Å map (map1). The final resolutions were established using Fourier Shell Correlation (FSC) at 0.143 cutoff (*56*) following gold-standard refinement. All conversions between Relion and CryoSPARC were performed using Daniel Asarnow’s pyem script (https://github.com/asarnow/pyem). The processing details and summaries are shown in figs. S3 and S4 and Table S1.

### Model building and refinement

Our cryo-EM reconstructions reveal BAP1/ASXL1 bound to nucleosome, with clearly resolved interfaces. The highest-resolution element in the reconstruction is the nucleosome core particle, akin to other published nucleosome-associated complexes(*19, 20*). BAP1/ASXL1 is very flexible; however, the quality of our maps allowed unambiguous rigid-body fitting and modeling of the proteins. To model the complex, we used a number of maps; the final BAP1/ASXL1-H2AK119Ub reconstruction from Titan Krios (Fig. S3 and S5) at 3.6 Å was used to describe the global architecture of the complex (map 1). The different maps used to build each region are summarized in Table S1. For the BAP1/ASXL1-H2AK119Ub complex, available X-ray crystal structures were used for initial rigid body fit into our cryo-EM maps. For the nucleosome, we used PDB IDs: 3tu4 (*36*), and 1ubq (*57*) for ubiquitin, and the structure of the homologous Calypso in complex with DEUBAD of ASX 6HGC (*19*) for BAP1 and ASXL1 respectively. The crystal structures were first manually fit into the density and then locally optimized using UCSF Chimera’s “Fit in map” function (*58*). Then we submitted the sequence of our constructs of BAP1, ASXL1 and Ub to AlphaFold-Multimer (*24*) and evaluated how the predicted models fit into our maps by performing a manual local optimization of the fitting of the best AlphaFold model using UCSF Chimera’s and the “Fit in map” function. We then used Coot (*24*) for local adjustments of secondary elements and side-chains into densities, using a combination of crystallographic and AlphaFold based models. The resulting complete model was refined in PHENIX (phenix.real_space_refine) (*59*) using secondary structure, ADPs, rotamer and Ramachandran restraints in 100 iterations. The model was then visually inspected, and Ramachandran outliers and problematic regions were fixed manually in Coot (final refinement statistics are summarized in Table S2). For assigning the residues interacting with the acidic patch and the DNA dyad of the nucleosome, we employed a second map with focused classification in these regions (map 2). An extra density could be observed near the N-terminal tail of BAP1 and the C-terminal tail of ASXL1, however, this region was too disordered in the cryo-EM reconstruction to reliably assign and model. To validate our structures, we first subjected the atoms to 0.1 Å displacement and then refined it in phenix.real_space_refine against one of the half-maps. This refined model was then converted to a 3D density map and compared against two half-maps and the summed map. We calculated FSC curves with half-map 1 (used for refinement, “work”, shown in our figures with red), half-map 2 (not participating in refinement, hence “free”, green) and the summed map (blue). We then tested the whole model against the 3.6 Å reconstruction (Fig S12). We only observed small differences between the ‘work’ and the ‘free’ FSC curves, which indicates lack of overfitting. Figures of the model and cryo-EM densities were prepared using Chimera, ChimeraX, Coot and PyMOL (*58, 60–62*).

### Deubiquitinase (DUB) catalytic activity assay

Deubiquitinase activity of wild-type BAP1/ASXL1 and indicated mutants towards the H2AK119ub1 dNuc substrate was assessed by monitoring fluorescence increase over 11 minutes in one-minute intervals at ambient temperature in the presence of Atto-532 labeled tUI free Ub sensor. 3.33 µL 3X DUB, 3.33 µL 3X H2AK119ub1 dNuc and 3.33 µL 3X tUI free Ub sensor (respectively: 5 nM, 10 nM, 10 nM final) in DUB buffer (20 mM HEPES pH 7.5, 20 mM NaCl, 2 mM MgCl2, 0.01% v/v Tween-20, 0.01% w/v BSA, and 10 mM DTT) were combined in a 384 well plate (PerkinElmer ProxiPlate-384 Plus F black, #6008260) and data gathered on a Envision 2150 plate reader (PerkinElmer) (excitation 531nm, emission 570, 555 mirror). The initial, linear portion of each duplicate DUB reaction was compared and data presented as Relative Fluorescence Units per minute (RFU/min). In the main text catalytic activity was also expressed as % of activity of each mutant relative to the WT. Deubiquitinase activity of wild-type BAP1/ASXL1 towards multiple ubiquinatied and unmodified nucleosome substrates was assessed as described above with the following modifications: the activity assay was using 2.5 nM DUB over eight minutes. The loss of catalytic activity was calculated as percent of activity of each mutant relative to the WT.

### Michaelis-Menten deubituitination kinetic assays (Michaelis–Menten curves)

Kinetic parameters were determined by monitoring BAP1/ASXL1 activity towards H2AK119ub1 dNuc substrate over the course of 11 minutes in one-minute intervals at ambient temperature. DUB and tUI free Ub sensor concentration were fixed while H2AK119ub1 dNuc concentration was varied in a 1.5 fold-dilution series. Final DUB concentrations were: wtBAP1/wtASXL1 at 0.5 nM; BAP1 R699A/R700A/R701A/wtASXL1 at 1 nM; wtBAP1/ASXl1 K243A/R244A/R246A at 1 nM; and BAP1 R56/R57/R59/R60A/wtASXL1 at 5 nM. For DUBs tested at 0.5 nM the substrate was varied from 300 to 3.5 nM; for DUBs tested at 1 nM from 300 to 5.2 nM; and for DUBs tested at 5 nM from 300 to 12 nM. In all cases, a no substrate sample was included for background correction. To calculate the amount of ubiquitin generated enzymatically, a free-ubiquitin standard curve (0.88 to 10 nM; 1.5-fold dilutions) was included, along with a no-ubiquitin sample for background correction. Three independent runs of triplicates were performed. Reaction volumes, tUI free Ub sensor concentration, assay plates and plate reader settings were as in the activity assay. H2AK119ub1 dNuc *K_0.5_*, *k_cat_*, and Hill coefficient (*h*) values were determined by applying an allosteric sigmoidal model using GraphPad 9.0 (Prism) to initial, linear velocities compared to H2AK119ub1 dNuc concentration. To determine initial velocities: first, a background correction was applied to the signal of the DUB samples and the standard curve. Second, a linear regression was applied to the background corrected standard curve at each timepoint. Third, the standard curve linear regression parameters were used to convert DUB background-corrected RFUs to the concentration of ubiquitin generated at each timepoint and a linear regression was fit to the data to generate initial velocity curves.

### Electro-Mobility Shift Assay (EMSA)

Assembled BAP1/ASXL1 complex was dialyzed into EMSA Buffer (10 mM HEPES pH 7.5, 50 mM NaCl, 0.1 mM DTT, 5% Glycerol). A 2-fold serial dilution (starting concentration between 1-5 µM) of BAP1/ASXL1 complex was made in EMSA Buffer. Non-hydrolyzable H2AK119Ub 187bp nucleosome was added to each reaction to reach a final concentration of 25 nM. Reactions were incubated on ice for 1 hour. 3.5 µL of reaction was analyzed on 4.5% native polyacrylamide gels (0.2 X TBE). Native acrylamide gels were stained with ethidium bromide and visualized using a Geldoc (Bio-Rad). The band quantification was performed with the program ImageLab (Bio-Rad). The amount of BAP1/ASXL1 bound to nucleosomes was determined by measuring the decrease in free nucleosome in each reaction using BAP1/ASXL1-free samples as background. The free DNA was taken under consideration for the calculation of free nucleosome. The apparent *K*d and the Hill coefficient for each binding curve were calculated by fitting the specific binding with Hill slope equation using the program Prism 9.0 (GraphPad). The final parameters were calculated using for at least 3 independent experiments (n:=3/data point). Data were plotted as mean ± s.e.

### Cell lines and cell culture

All mESC lines were grown on 0.1% gelatin-coated dishes in 2i/LIF-containing GMEM medium (Euroclone) supplemented with 16% fetal calf serum (Euroclone), 2 mM glutamine (GIBCO), 100 U/mL penicillin, 0.1 mg/mL streptomycin (GIBCO), 0.1 mM non-essential amino acids (GIBCO), 1 mM sodium pyruvate (GIBCO), 50 μM β-mercaptoethanol phosphate buffered saline (PBS; GIBCO), 1000 U/mL leukemia inhibitory factor (LIF; produced in-house), and GSK3β and MEK 1/2 inhibitors (ABCR GmbH) to a final concentration of 3 μM and 1 μM, respectively.

Stable expressing clones BAP1 KO ESC were generated by transfecting 10 µg pCAG expression vectors encoding N-terminal 2xFlag-HA-tagged BAP1 wild-type or with C91S; R56E/R59E; R699E/R700E; R699A/R700A/R701A; and R56A/R57A/R59A/R60A mutations using Lipofectamine 2000 (ThermoFisher Scientific), as per manufacturer’s instructions. 24 h post-transfection ESC were selected with neomycin (0,5 μg/mL) for 5 days. Cells were then split to clonal density (∼1:50) onto 15cm plates. Clones were isolated 12 days later and grown further before screening for rescue allele expression by Western blot.

### Western Blot

Western blot analyses were performed with total protein lysates. All mESCs lines were lysed and sonicated in ice-cold S300 buffer (20 mM Tris-HCl pH 8.0, 300 mM NaCl, 10% glycerol, 0.2% NP40) and supplemented with Benzonase (25U/μL Millipore) and protease inhibitors (Roche). Precipitates were removed by centrifugation. Clear lysates were resuspended in Laemmli sample buffer and boiled for 5 minutes. Protein lysates were separated on SDS-PAGE gels and transferred to nitrocellulose membranes. After probing with the appropriate primary and secondary antibodies, chemiluminescence signals were captured with the ChemiDoc Imaging System (Bio-Rad). Western blots were performed with: anti-BAP1 (D7W70; CST), anti-H2AK119ub1 (8240; CST), anti-H3 (ab1791; Abcam).

### Quantification and statistical analysis

Protein quantification was done using an A280 extinction coefficient of 106,660 M^−1^cm^−1^ for BAP1/ASXL1 complex on a Nanodrop spectrophotometer (Thermo-Fisher).

## Supplementary Text

### Extended Figure Legends

**Figure 3A:** Residues that mediate interaction of BAP1/ASXL1 with the H2AK119Ub nucleosome were assessed for their contribution to deubiquitinase (DUB) activity against a H2AK119ub1 dNuc substrate. Activity of mutations to BAP1 catalytic site (C91S), acid patch interaction residues (BAP1 R56A/R57A/R59A/R60A; or BAP1 R56E/R59E), DNA dyad interaction residues (BAP1 R699D/R700D; BAP1 R699A/R700A/R701A; or ASXL1 K243A/R244A/R246A) or DNA exit residues (ASXL1 R336A/R338A/R343A/R346A) toward H2AK119ub1 dNuc were compared to WT BAP1/ASXL1 *in vitro*. A fluorescently labeled tUI free Ub sensor (a ubiquitin binding domain containing protein that undergoes a fluorescence change when it engages free ubiquitin) was used to monitor ubiquitin release over 11 minutes at ambient temperature in duplicate (5 nM DUB, 10 nM H2AK119ub1 dNuc substrate, 10 nM tUI free Ub sensor) and the initial, linear reaction rates presented as Relative Fluorescence Units per minute (RFU/min). When free ubiquitin is released from the Kub1-dNuc by the DUB, it binds to the tUI free Ub sensor, resulting in a fluorescence increase. Comparison of initial velocities for each mutant complex relative to WT reveals the extent of activity impairment.

**Figure 3H & I:** tUI free Ub sensor was used to monitor WT BAP1/ASXL1; BAP1 R699A/R700A/701A/wtASXL1; wtBAP1/ASXL1 K243A/R244A/R246A [**Fig3H**, DNA dyad interaction mutants] and BAP1 R56A/R57A/R59A/R60A/wtASXL1 [**Fig3I**, acidic patch interaction mutant] activity over 11 minutes at ambient temperature. DUB (0.5 nM, 1 nM, 1 nM, 5 nM respectively) and sensor (10 nM) concentration was fixed while varying the H2AK119ub1 dNuc substrate. A free Ub standard curve was used to calculate the amount of enzymatically generated ubiquitin, with initial velocities relative to H2AK119ub1 concentration used to determine H2AK119ub1 dNuc *K*_0.5_, *k*_cat_, and Hill coefficient values by applying an allosteric sigmoidal model. Representative results of three independent experiments (performed in triplicate) are displayed. As compared to WT BAP1/ASXL1, BAP1 R699A/R700A/R701A/wtASXL1 (DNA dyad) [H] impacts binding affinity whereas wtBAP1/ASXL1 K243A/R244A/R246A (DNA dyad) [H] and BAP1 R56A/R57A/R59A/R60A/wtASXL1 (acidic patch) [I] impact catalytic turnover.

**Fig. S1.**
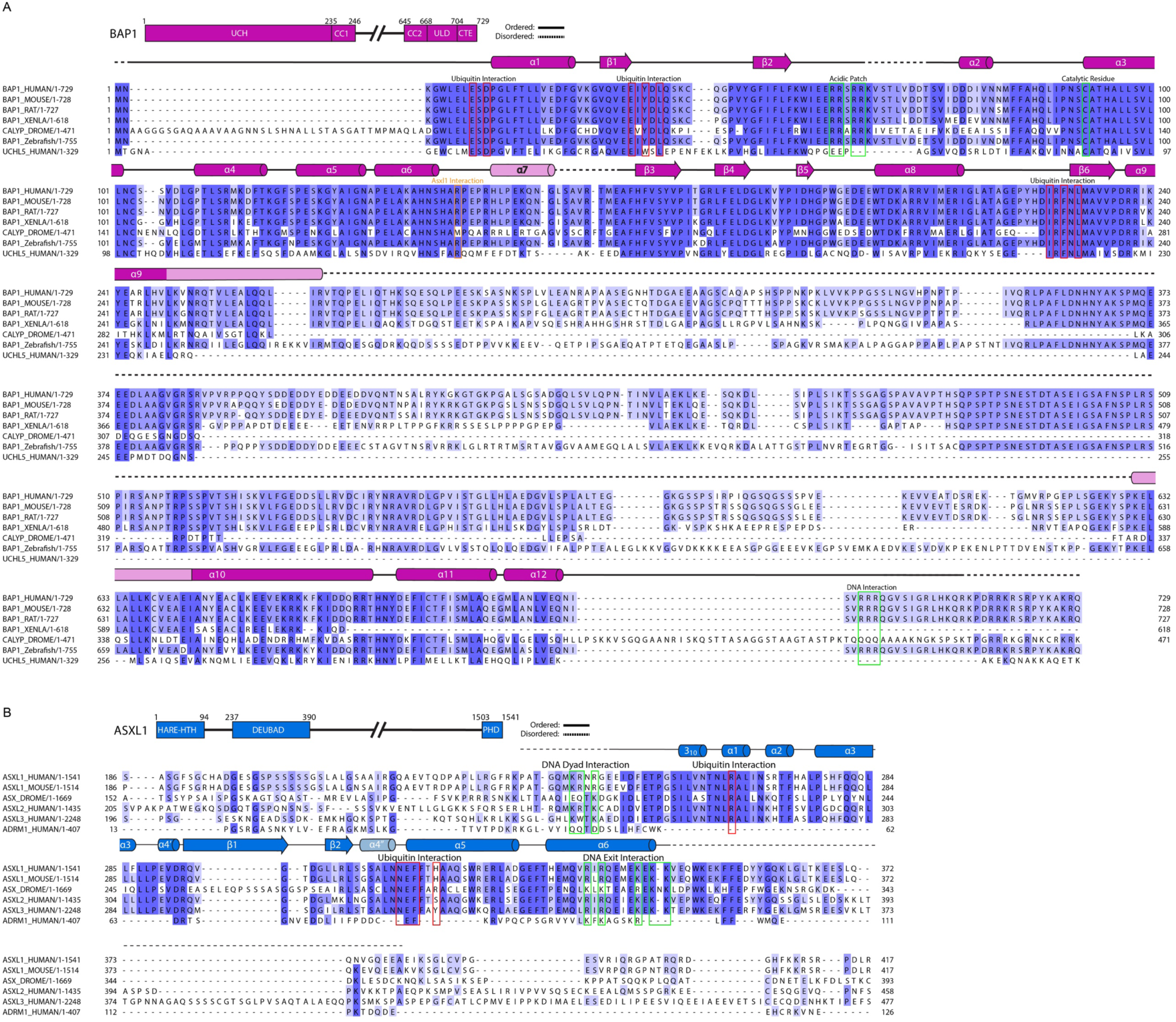
BAP1 and ASXL1 sequence alignment. (Top) Protein alignment using MUSCLE to align BAP1, Calypso and UCH-L5 from different organisms. Secondary structure elements present in our structure are shown above sequence alignment. Residues of BAP1 forming the interfaces with Ub, acidic patch, DNA dyad, DNA exit, and the catalytic residue are boxed and labeled. (Bottom) Protein alignment using MUSCLE to align ASXL1 homologs across species (human, mouse, *Drosophila*) and within human cells (ASXL1, ASXL2. ASXL3, RPN13 (ADRM1)). Secondary structure elements in our cryo-EM structure are shown above sequence alignment. Residues of ASXL1 forming the interfaces with Ub, acidic patch, and DNA exit are boxed and labeled.

**Fig. S2.**
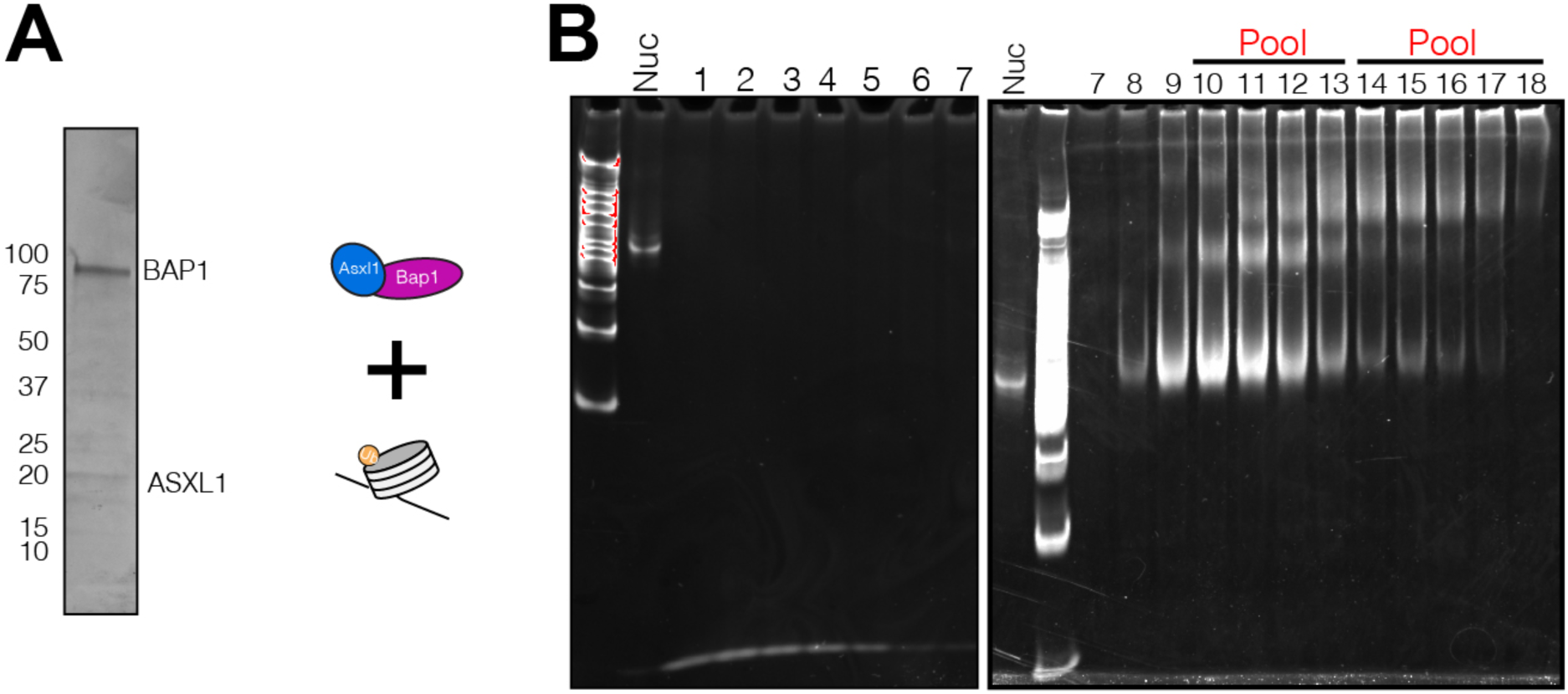
BAP1/ASXL1-H2AK119Ub nucleosome sample preparation for structural analysis. A) Representative BAP1/ASXL1 complex analyzed by SDS-PAGE (Left). Graphical depiction of BAP1/ASXL1-H2AK119Ub nucleosome assembly (Right). B) Native PAGE analysis of GraFix fractions from BAP1/ASXL1-H2AK119Ub nucleosome assembly. Gradient fractions pooled and used in Cryo-EM are indicated.

**Fig. S3.**
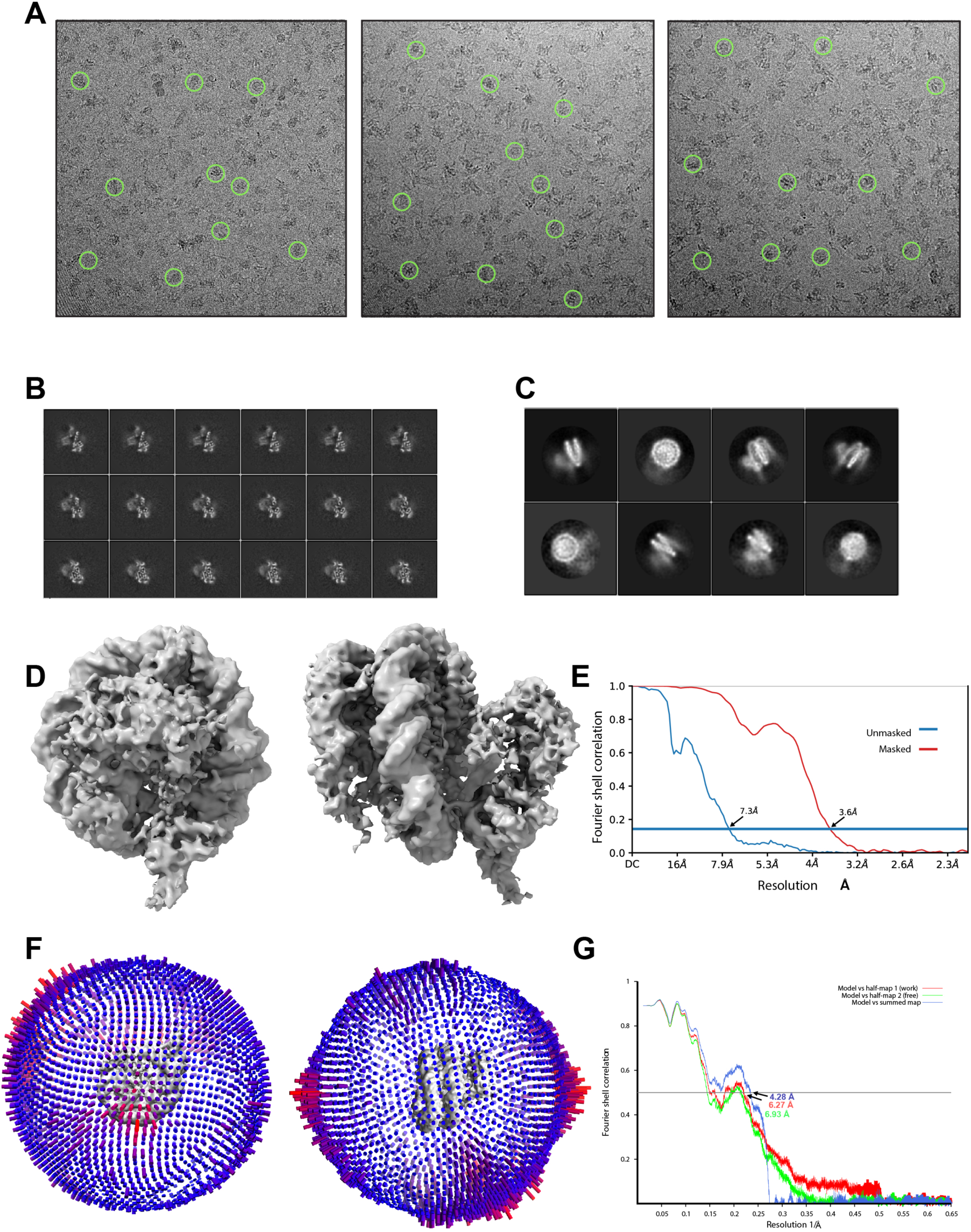
Cryo-EM analysis of the BAP1/ASXL1-H2AK119Ub nucleosome complex. A) Raw cryo-EM images of BAP1/ASXL1-H2AK119Ub complex were collected as described in Materials and Methods. Representative micrographs from the collection with selected particles of the complex are shown in green circles. B) Slices at different levels along the Z-axis through the final cryo-EM map. C) Representative 2D class averages selected from the dataset. D) Two representative views of the 3D reconstruction of BAP1/ASXL1–H2AK119Ub nucleosome complex. E) FSC plot of the 3.6 Å BAP1/ASXL1–H2AK119Ub complex between two independently refined half maps (measured at FSC=0.143). F) Euler angle distribution of assignment of particles used to generate the final 3D reconstruction of the 3.6 Å complex. The length of every cylinder is proportional to the number of particles assigned to the specific orientation. G) Cross-validation of the model built against the cryo-EM reconstruction (see Materials and Methods): FSC curves for the model and cryo-EM map calculated from the final model and half map 1 (‘work’, red), half map 2 (‘free’, green) and summed map (blue).

**Fig. S4.**
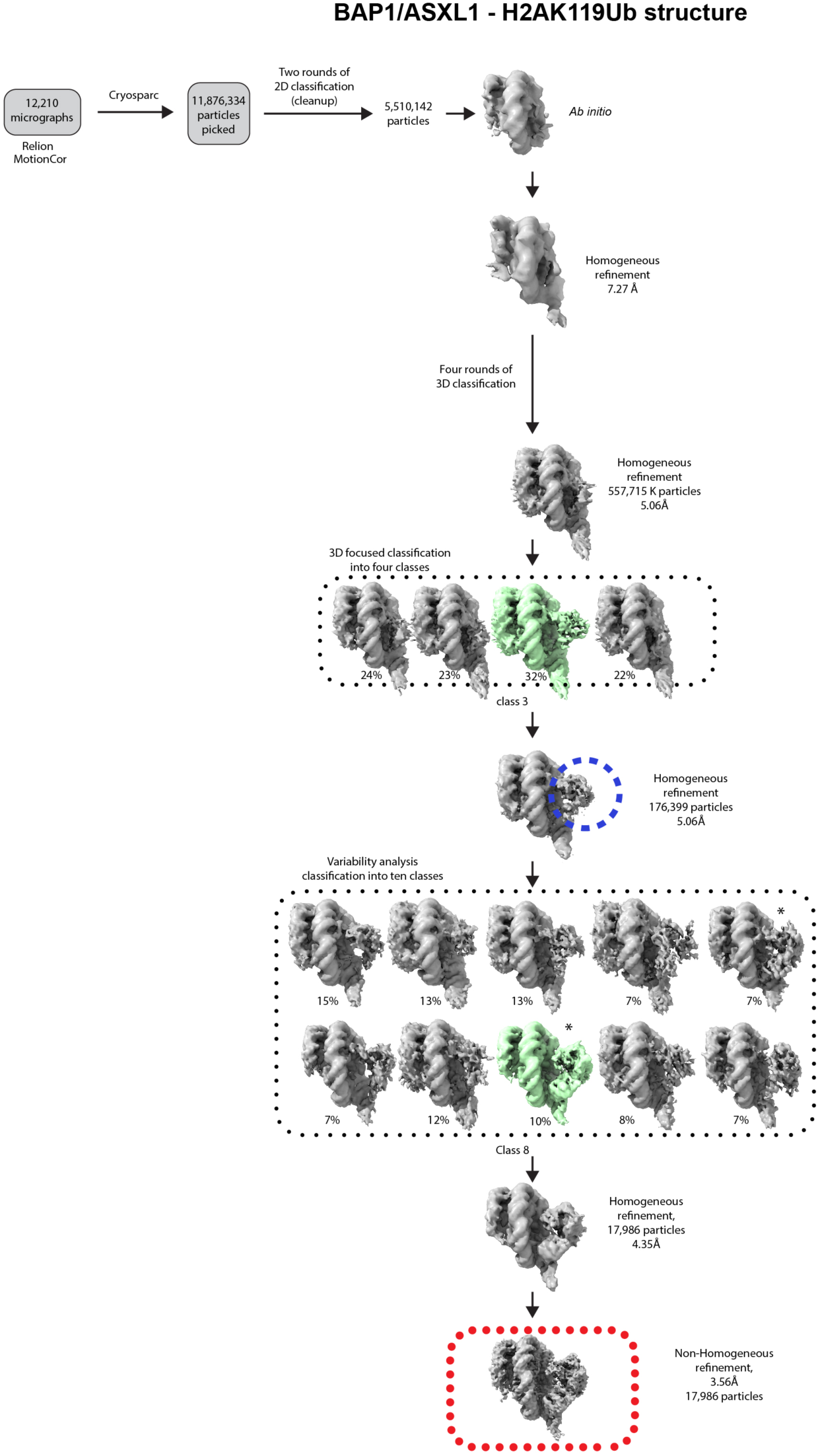
3D classification scheme of BAP1/ASXL1–H2AK119Ub nucleosome samples. Processing scheme for classification of BAP1/ASXL1-H2AK119Ub nucleosome complex dataset collected on Titan Krios operated at 300 kV. Images represent the path that led to the final subset, with particles and map selected for each step shown in green. A mask applied to BAP1/AXSL1 in 3D focused classification and variability analysis is shown in a blue circle. In the variability analysis step, the most stable conformation (classes 5 and 8) is marked with an asterisk (*), with the remaining 3D classes in a more flexible second conformation. Superposition of classes 6 and 8 of the variability analysis step is shown in Fig. S12

**Fig. S5.**
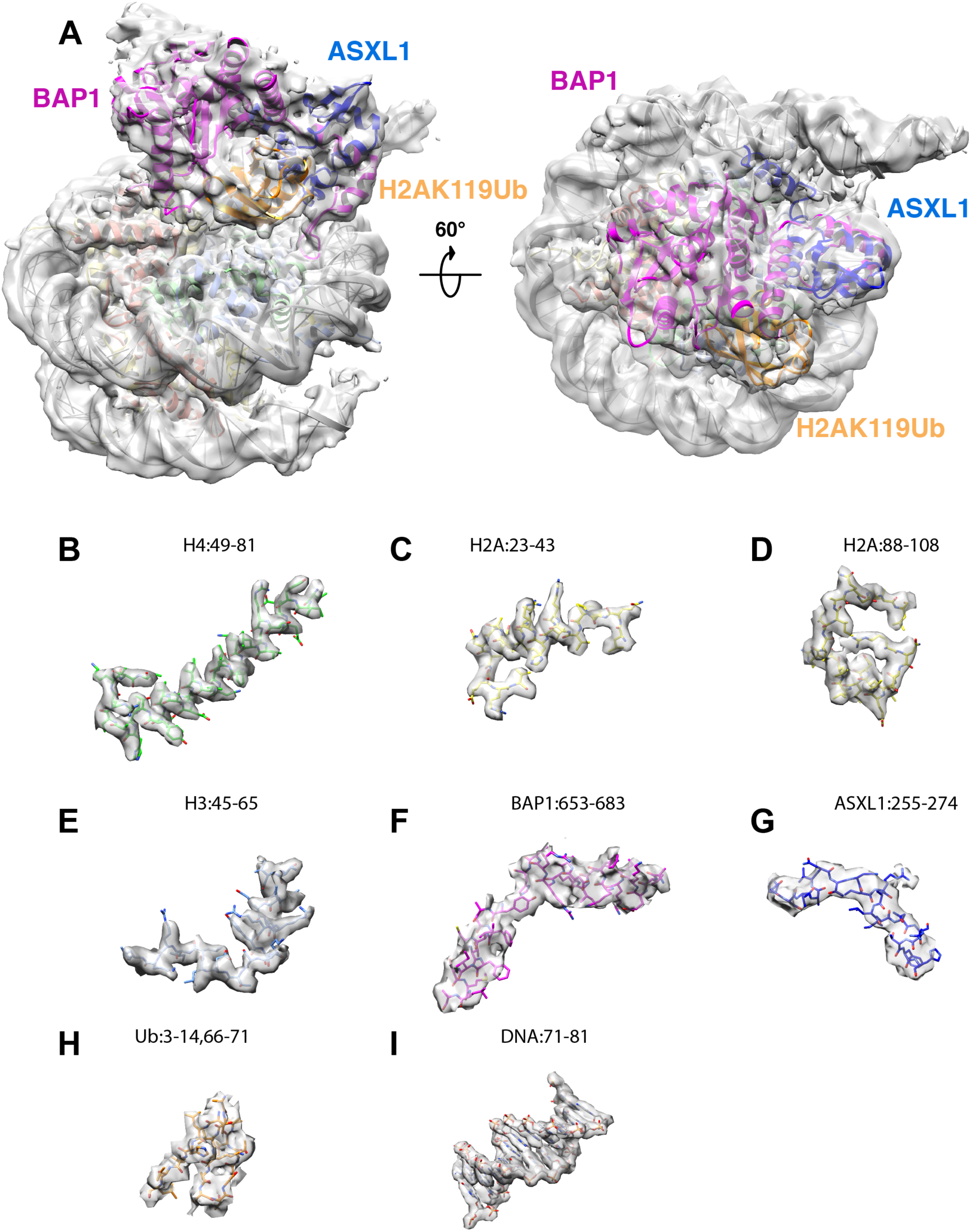
Different regions of the cryo-EM map of BAP1/ASXL1-H2A119KUb nucleosome complex. Selected views of the model fit to the cryo-EM map for the BAP1/ASXL1-H2AK119Ub nucleosome structure are shown. A) Overview of the complete cryo-EM map/model. B)-I) Selected regions of the model fit to cryo-EM map are shown. A close-up of the model/cryo-EM map fitting in the regions of interaction with the acidic patch and DNA clamp is shown in Fig. S8. The maps were visualized with Chimera.

**Fig. S6.**
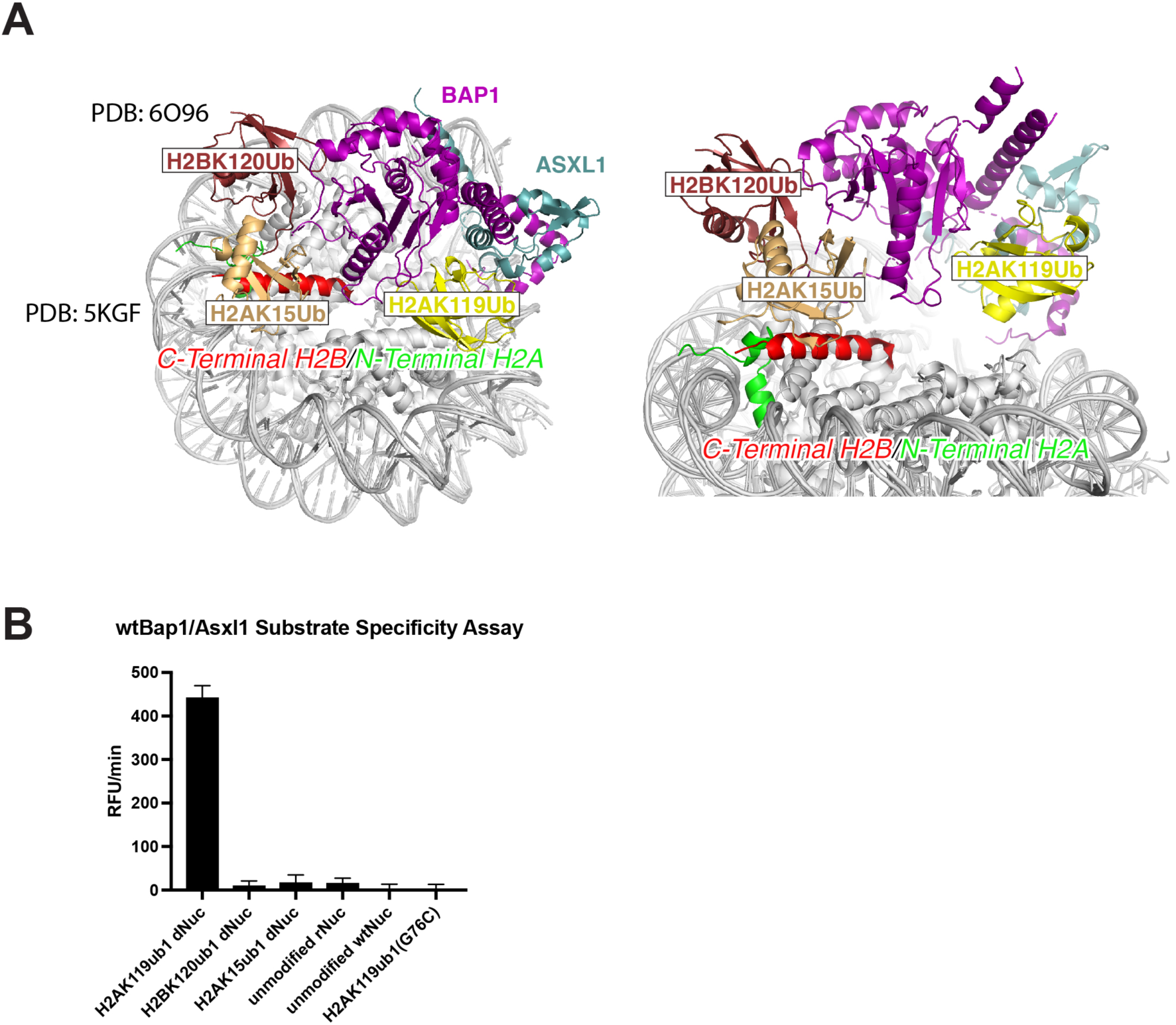
Structural alignment and activity assays showing incompatibility of BAP1/ASXL1 with nucleosomal H2BK120Ub and H2AK15Ub. A) Our cryo-EM structure was aligned to structures of chromatin modifiers in complex with H2BK120Ub (PDB ID 6o96) (*31*)and H2AK15Ub (PDB 5kgf) (*63*) nucleosomes. The positions of Ub in both complexes are shown in superposition to our BAP1/ASXL1-H2AK119Ub nucleosome complex. H2BK120Ub, H2AK15Ub, and H2AK119Ub are shown and labeled. B) WT BAP1/ASXL1 dNuc substrate specificity was assessed *in vitro*, comparing H2AK119ub1, H2BK120ub1, H2AK15ub1, unmodified rNuc, unmodified wt nucleosome and crosslinked, nonhydrolyzable nucleosome. BAP1/ASXL1 activity was monitored using the tUI free Ub sensor over 8 minutes at ambient temperature in duplicate (2.5 nM DUB, 10 nM dNuc, 10 nM tUI free Ub sensor) and initial, linear reaction rates are presented as Relative Fluorescence Units per minute (RFU/min). Free ubiquitin released by BAP1/ASXL1 binds the tUI free Ub sensor, leading to an increase in fluorescence. WT BAP1/ASXL1 is only able to use H2AK119ub1 dNuc as a substrate.

**Fig. S7.**
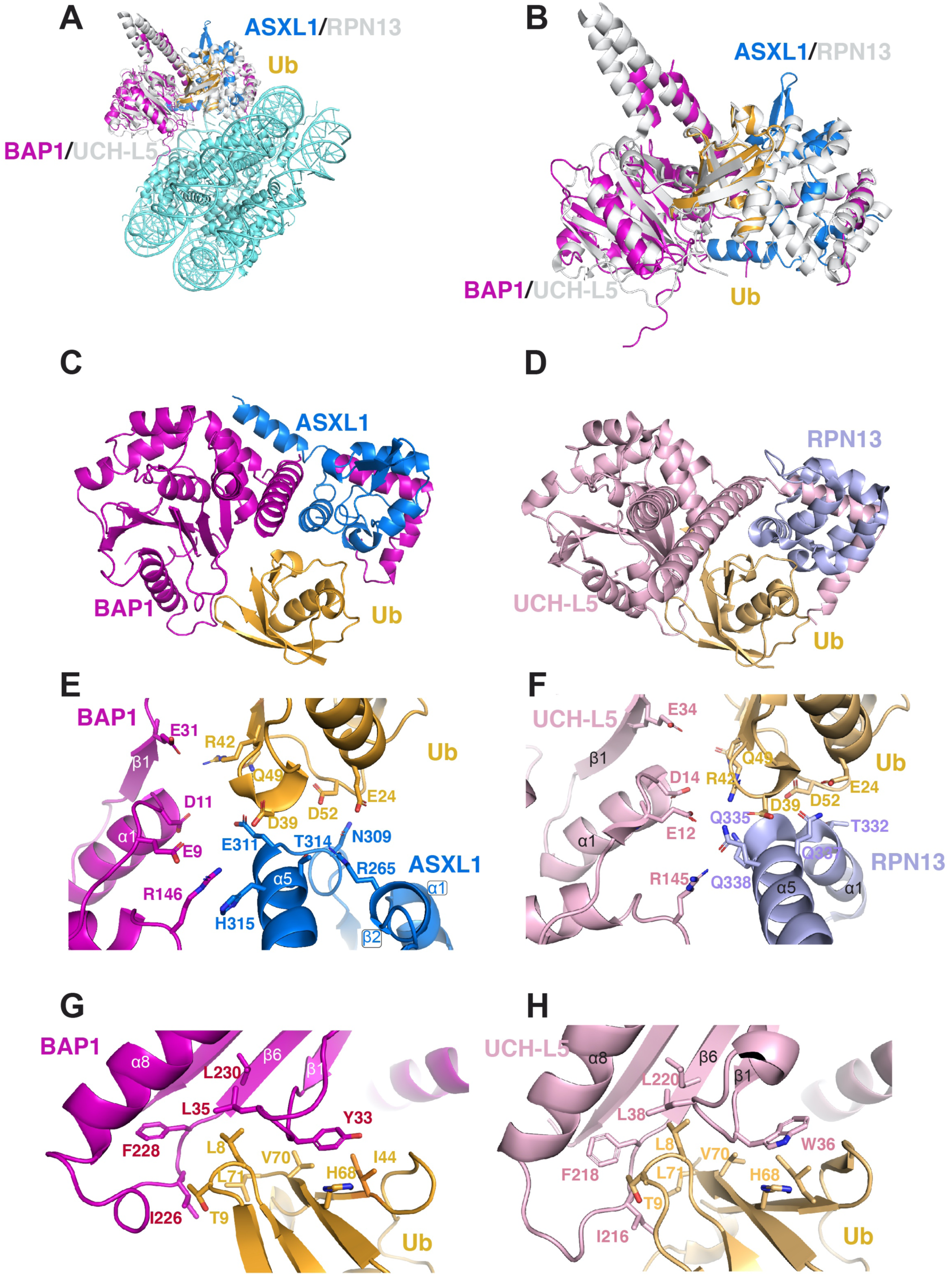
The interfaces of BAP1/ASXL1 and UCHL5/RPN13 with Ub are conserved. A) Superposition between BAP1/ASXL1-H2AK119Ub nucleosome complex determined in this work and UCH-L5/RPN13 (PDB ID 4uel)(*21*), B) Close up of A) showing the Ub interaction of each complex. Side-by-side comparison showing Ub interaction with C) BAP1/ASXL1 and D) UCH-L5/RPN13. Side-by-side comparison of the electrostatic interactions with Ub in the interface of E) ASXL1-BAP1 and F) RPN13-(UCH-L5). Side-by-side comparison of the hydrophobic interactions with Ub in the interface of G) BAP1 and H) UCH-L5. Proteins are color coded as Fig. 2.

**Fig. S8.**
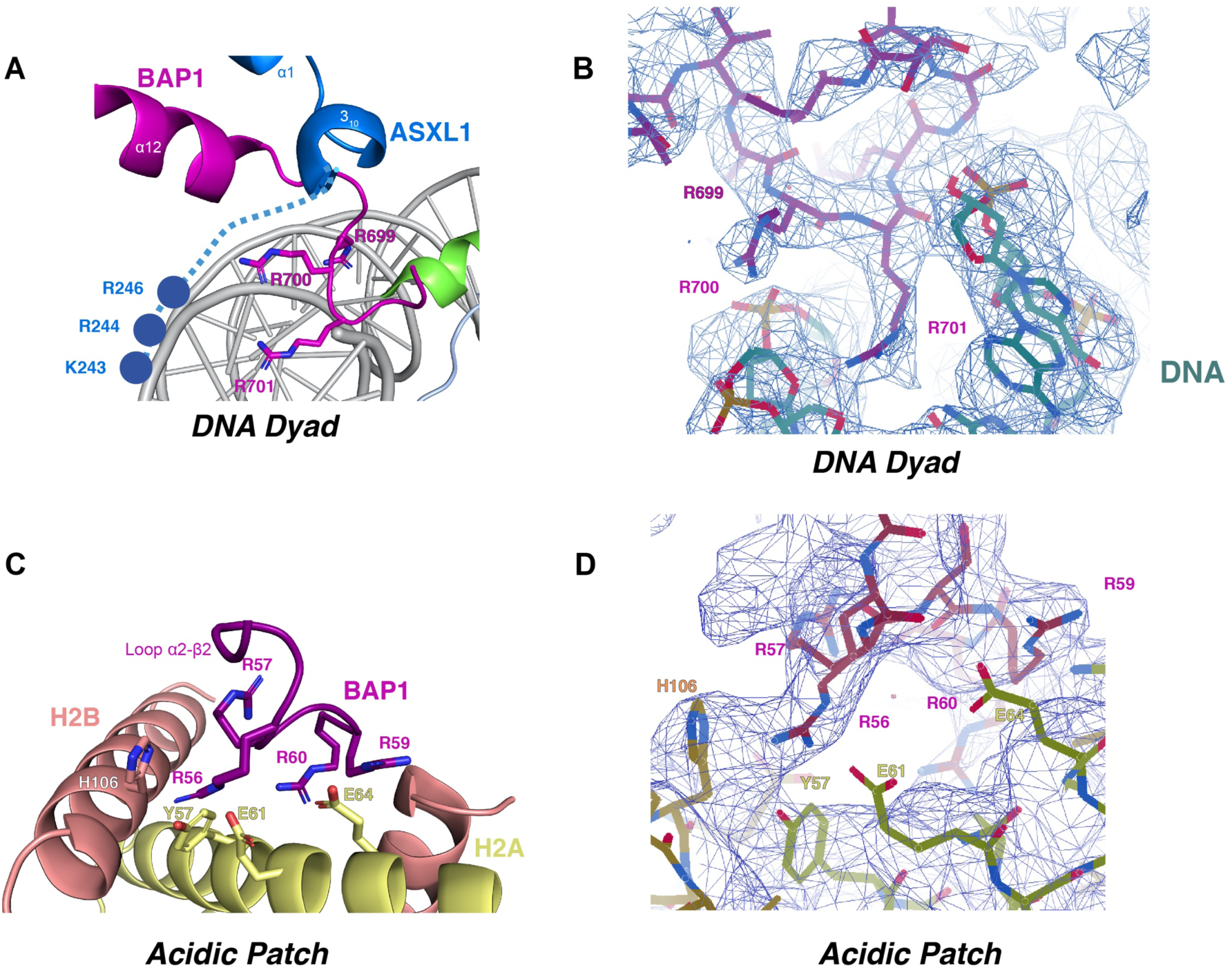
Different regions of the cryo-EM map of BAP1/ASXL1-H2A119KUb nucleosome complex. Selected views of the model fit to the cryo-EM maps for the structure are shown. A) DNA Clamp model. B) Map for the region in A (map 3). C) Acidic patch interface. D) Map for the region in C (map 2). The maps were visualized with Coot.

**Fig. S9.**
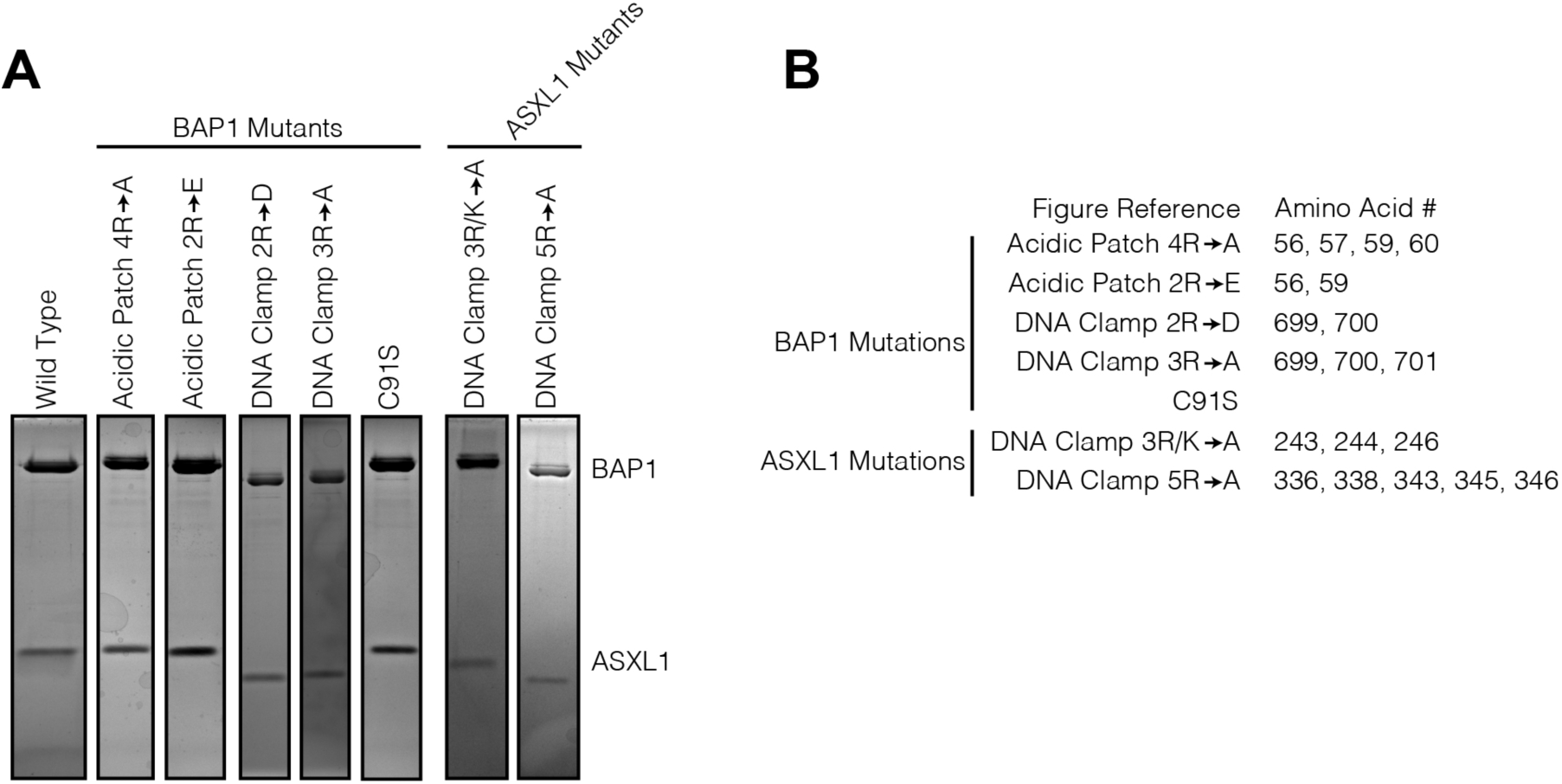
BAP1/ASXL1 complex assembly for biochemical characterization. A) BAP1/ASXL1 complex analyzed by SDS-PAGE after size exclusion chromatography. B) Further information regarding BAP1/ASXL1 mutations in panel A.

**Fig. S10.**
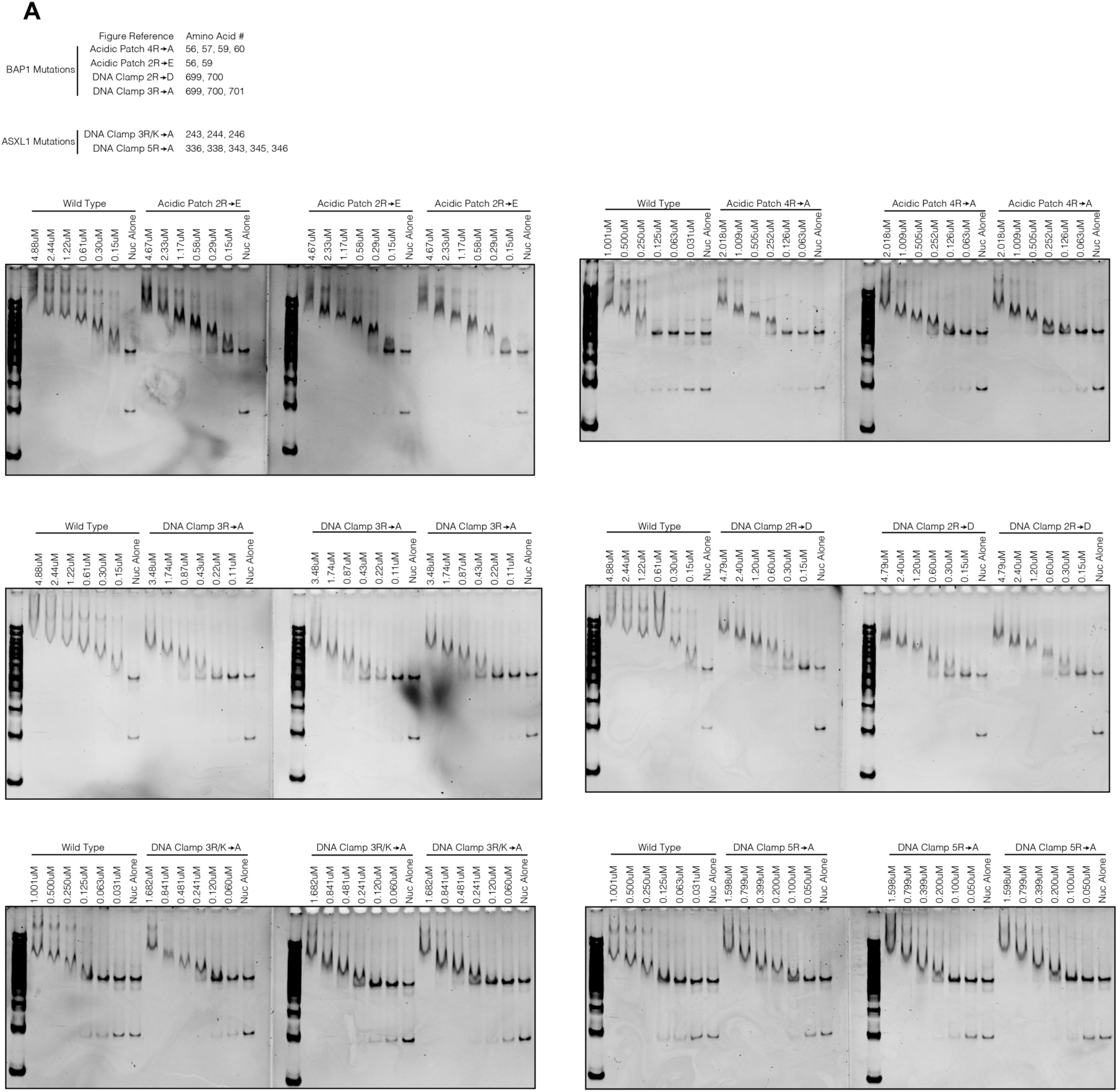
Raw data for EMSA of BAP1/ASXL1 mutants. A) Information about BAP1/ASXL1 mutations (top left). Native PAGE of BAP1/ASXL1 bound to H2AK119Ub nucleosomes. BAP1/ASXL1 mutation status and concentration, are marked on gels.

**Fig. S11.**
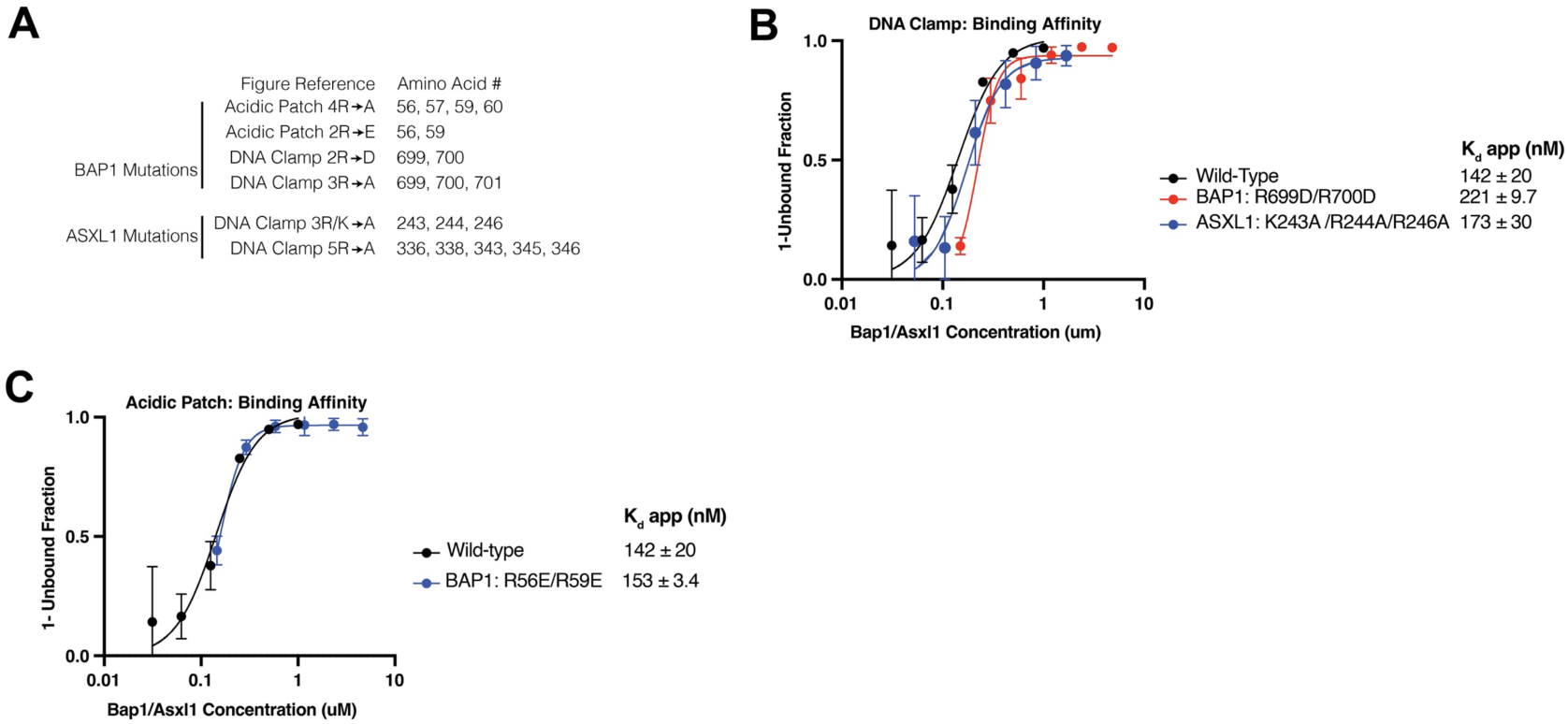
Biochemical characterization by EMSA and catalytic activity assay. A) Summary of information of BAP1/ASXL1 mutations is shown. Nucleosome binding curves for the B) DNA clamp mutants and C) BAP1 acidic patch interaction mutant, measured by EMSA. Each data point and error bar indicate the mean ± SD from three independent experiments. The standard errors of dissociation constants (Kd) are indicated.

**Fig. S12.**
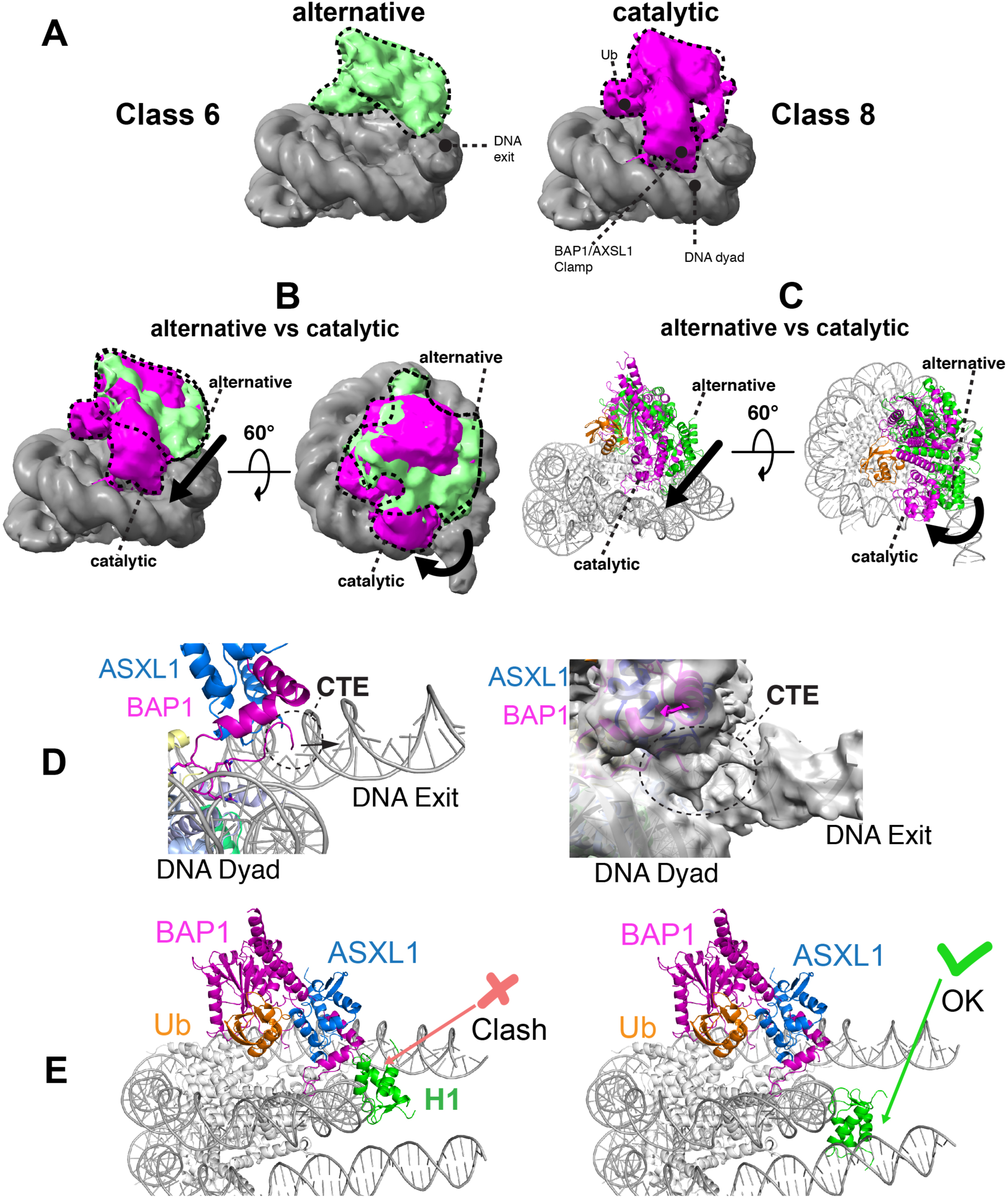
Conformations of BAP1/ASXL1 on the nucleosome. A) Two different cryo-EM maps from variability analysis (shown in Fig. S4) depicting an alternative, more flexible conformation in class 6 (left) compared with the more stable conformation of BAP1/ASXL1-H2AK119Ub nucleosome in class 8 (right), and B) superposition of both classes. (C) Model from superposition shown in (B). The more stable conformation is characterized by the binding of the BAP1/ASXL1 clamp near the DNA dyad. BAP1/ASXL1 clamp might bind to the DNA exit rather than the DNA dyad in the more flexible conformation. Maps filtered with Gaussian filter at 2.5 SD. D) BAP1 CTE region observed in our structure shown in the model (left) and the fitting to an unsharpened (map 1). E) Superposition of H1 in the structure of BAP1/ASXL1-H2AK119Ub nucleosome determined in this study, with the structure of the chromatosome (PDB ID 7pfv)(*43*), showing the (left) incompatibility between linker histone H1.4 (when it is bound to DNA at the side of BAP1/ASXL1) and BAP1/ASXL1 clamp at the dyad axis, also near the usual conformation of the H2A docking domain; and (right) the compatibility of BAP1/ASXL1 when H1.4 is bound on the other side of the DNA linker.

**Fig. S13.**
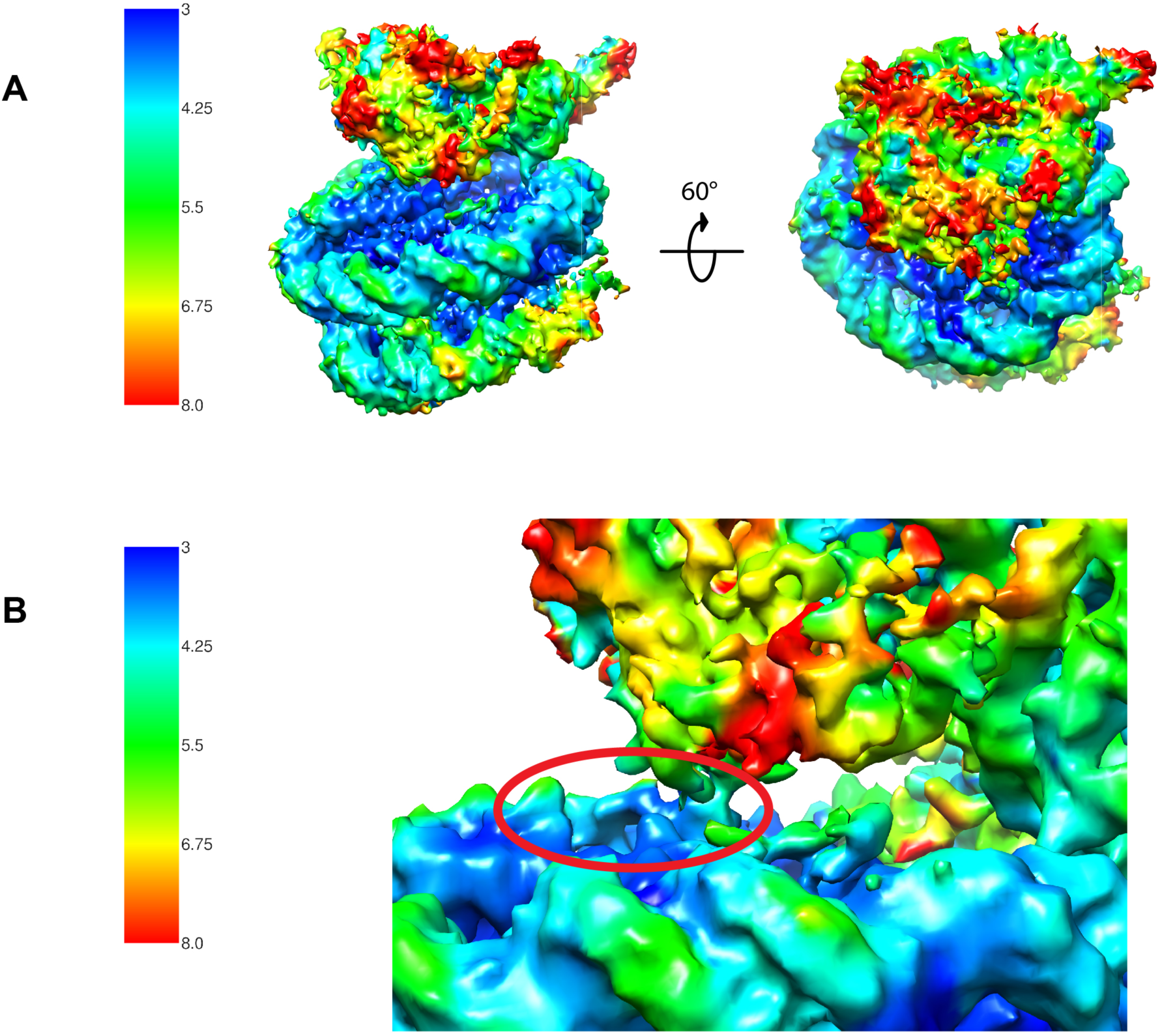
Local resolution heat map for the reconstruction for BAP1/ASXL1-H2AK119Ub nucleosome complex. The local resolution heat map (of map 1), calculated using cryoSPARC’s built-in local resolution algorithms, is shown in A) (left) Front view of the Coulomb potential maps visualized in Chimera (right). B) Close-up of the R-finger region of BAP1 interacting with the acidic patch. The model fit to the cryo-EM map in this region is shown in Fig S8.

**Fig. S14.**
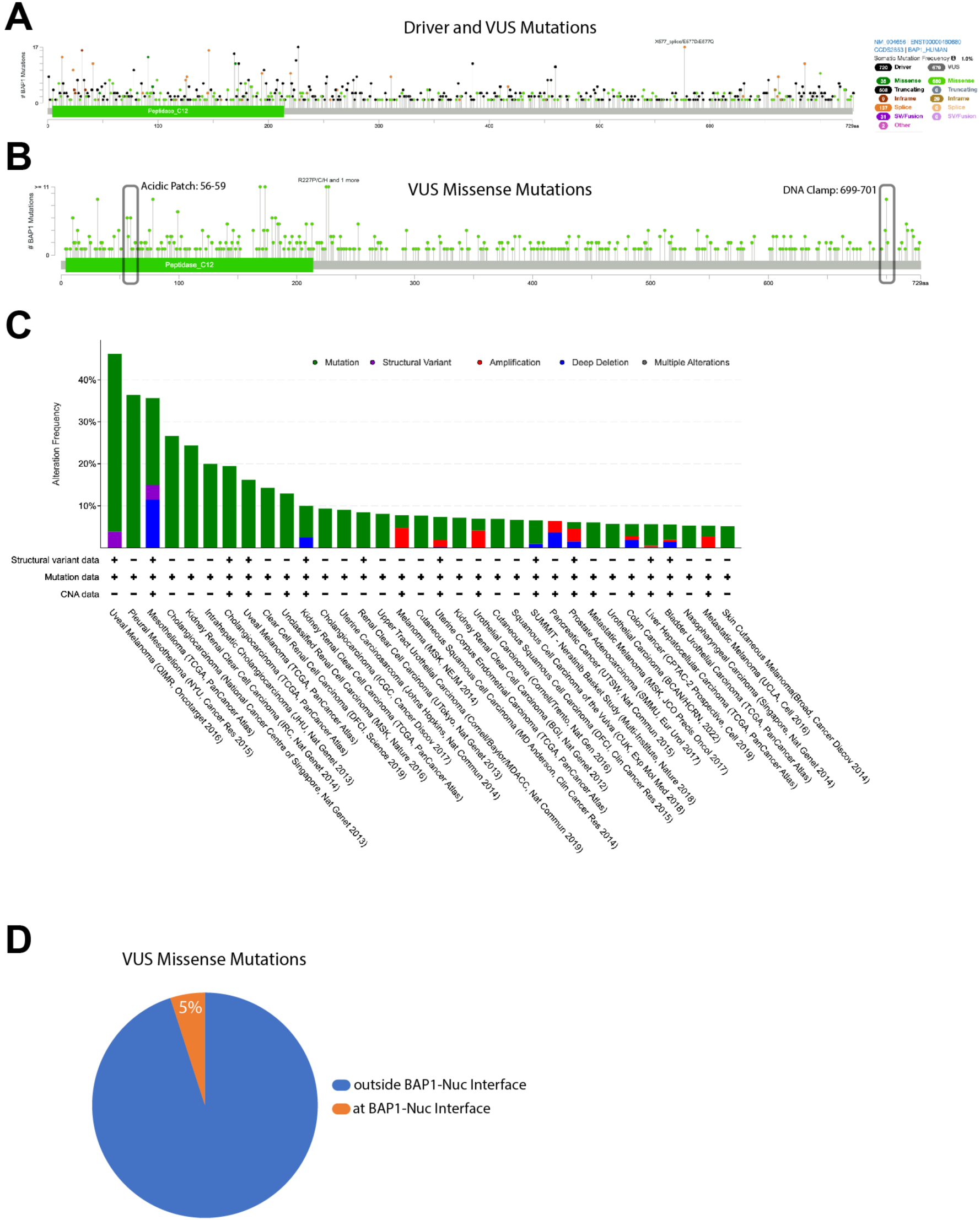
BAP1 mutations found in cancer. Searching the cBioPortal database for a curated set of non-redundant cancer studies on BAP1. A and B) Driver and VUS mutations are mapped across the amino acid sequence of BAP1 with mutational frequency represented as high. C) BAP1 cancer associated mutations compared with alteration frequency and associated disease. D) Pie chart representation of the total number of VUS mutations and the number of VUS mutations at BAP1-Nucleosome interfaces.

**Fig. S15.**
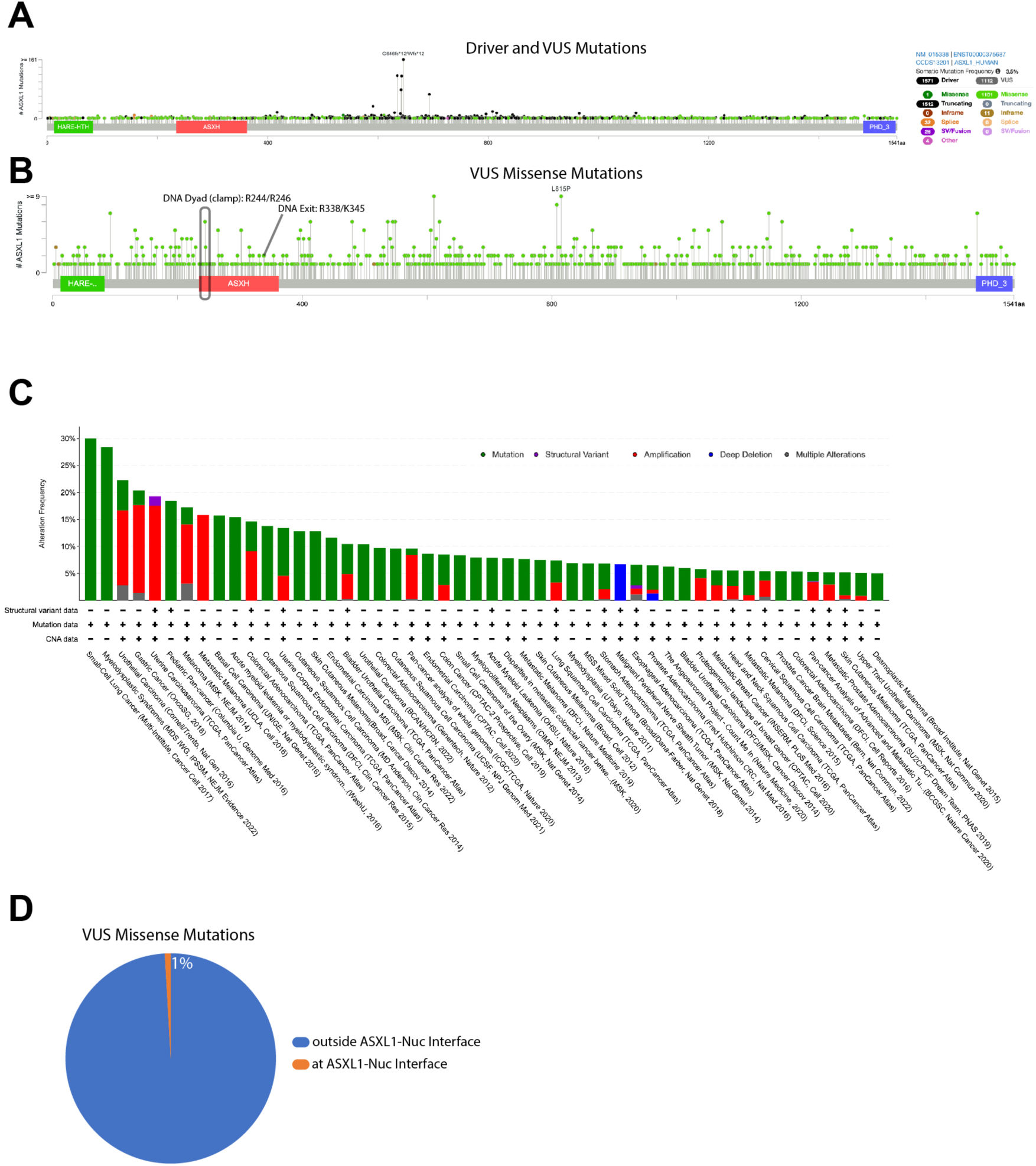
ASXL1 mutations found in cancer. Searching the cBioPortal database for a curated set of non-redundant cancer studies on ASXL1. A and B) Driver and VUS mutations are mapped across the amino acid sequence of ASXL1 with mutational frequency represented as high. C) ASXL1 cancer associated mutations compared with alteration frequency and associated disease. D) Pie chart representation of the total number of VUS mutations and the number of VUS mutations at BAP1-Nucleosome interfaces.

**Table S1.**
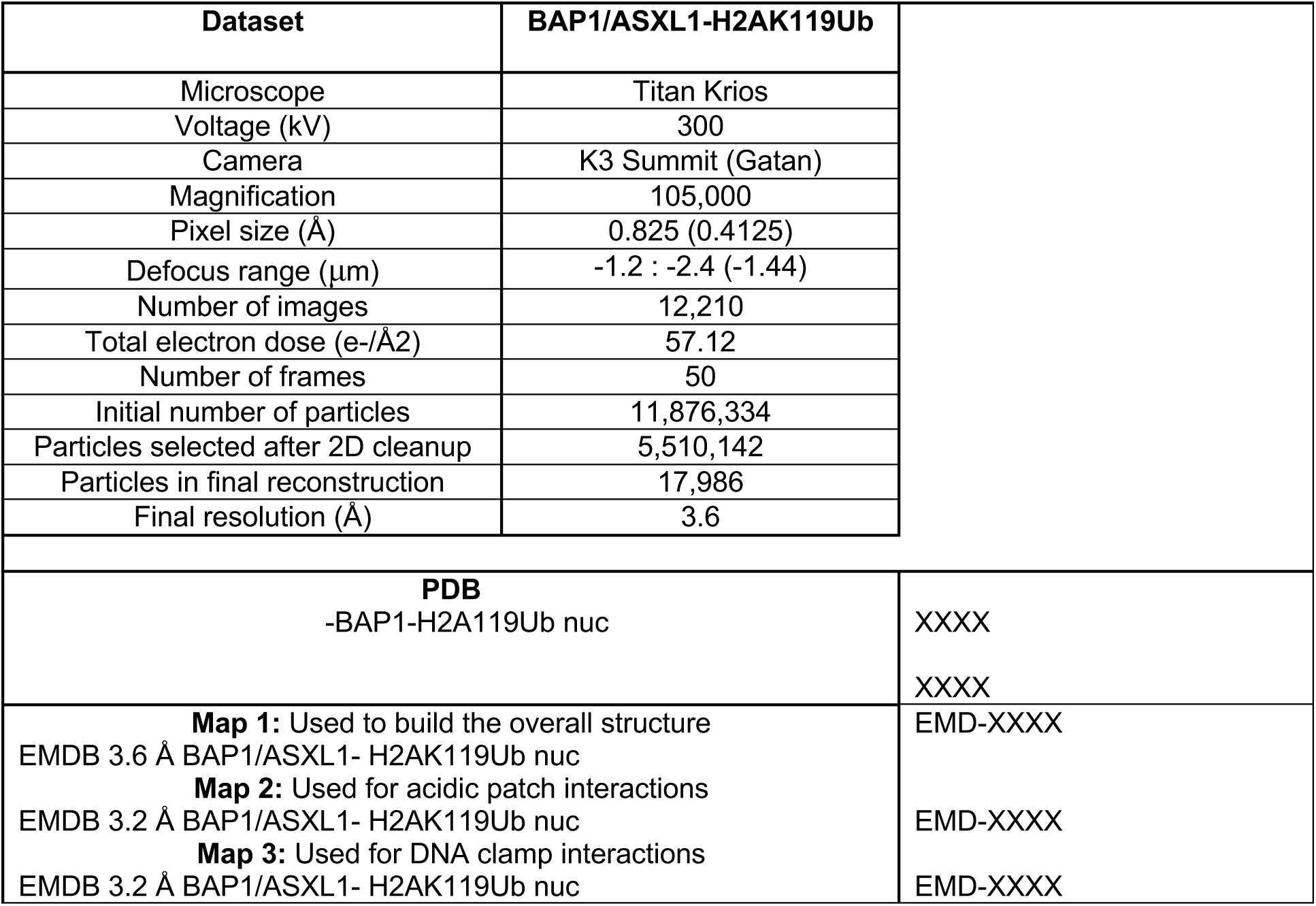
Summary for cryo-EM data collection, refinement and deposition.

| Dataset | BAP1/ASXL1-H2AK119Ub |  |
| --- | --- | --- |
| Microscope | Titan Krios |  |
| Voltage (kV) | 300 |  |
| Camera | K3 Summit (Gatan) |  |
| Magnification | 105,000 |  |
| Pixel size (Å) | 0.825 (0.4125) |  |
| Defocus range (µm) | -1.2 : -2.4 (-1.44) |  |
| Number of images | 12,210 |  |
| Total electron dose (e-/Å<sup>2</sup>) | 57.12 |  |
| Number of frames | 50 |  |
| Initial number of particles | 11,876,334 |  |
| Particles selected after 2D cleanup | 5,510,142 |  |
| Particles in final reconstruction | 17,986 |  |
| Final resolution (Å) | 3.6 |  |
| **PDB**<br>-BAP1-H2A119Ub nuc |  | XXXX |
|  |  | XXXX |
| **Map 1:** Used to build the overall structure<br>EMDB 3.6 Å BAP1/ASXL1- H2AK119Ub nuc<br>**Map 2:** Used for acidic patch interactions<br>EMDB 3.2 Å BAP1/ASXL1- H2AK119Ub nuc<br>**Map 3:** Used for DNA clamp interactions<br>EMDB 3.2 Å BAP1/ASXL1- H2AK119Ub nuc |  | EMD-XXXX<br>EMD-XXXX<br>EMD-XXXX |

**Table S2.** Summary for Cryo-EM data collection, refinement and validation statistics.

|  |  |
| --- | --- |
|  | BAP1/ASXL1-H2A119Ub nucleosome (EMD-XXXXX) (PDB NXXX) |
| **Data collection and processing** |  |
| Magnification | 105,000x |
| Voltage (kV) | 300 |
| Electron exposure (e-/Å<sup>2</sup>) | 57.12 |
| Defocus range (µm) | -1.2 to -2.4 (-1.44) |
| Pixel size (Å) | 0.825 (0.4125) |
| Symmetry imposed | *C1* |
| Initial particle images (no.) | 11,876,334 |
| Final particle images (no.) | 17,986 |
| Map resolution (Å) | 3.6 |
| 0.143 FSC threshold |  |
| Map resolution range (Å) | 3.6-11 |
| **Refinement** |  |
| Initial model used (PDB code) | 3tu4, 6hgc, 1ubq, AlphaFold |
| Model resolution (Å) | 3.6 |
| FSC threshold (0.143) |  |
| Model resolution range (Å) | 3.6-11 |
| Map sharpening *B* factor (Å<sup>2</sup>) | -70 |
| Model composition |  |
| Non-hydrogen atoms | 15,911 |
| Protein residues | 1208 |
| Nucleotides | 308 |
| *B* factors (Å<sup>2</sup>) |  |
| Protein | 78.22 |
| Nucleotides | 139.38 |
| R.m.s. deviations |  |
| Bond lengths (Å) | 0.009 |
| Bond angles (°) | 1.144 |
| Validation |  |
| MolProbity score | 2.0 |
| Clashscore | 9 |
| Poor rotamers (%) | 1 |
| Ramachandran plot |  |
| Favored (%) | 96.41 |
| Allowed (%) | 3.58 |
| Disallowed (%) | 0.0 |

**Supplementary Table S3.**
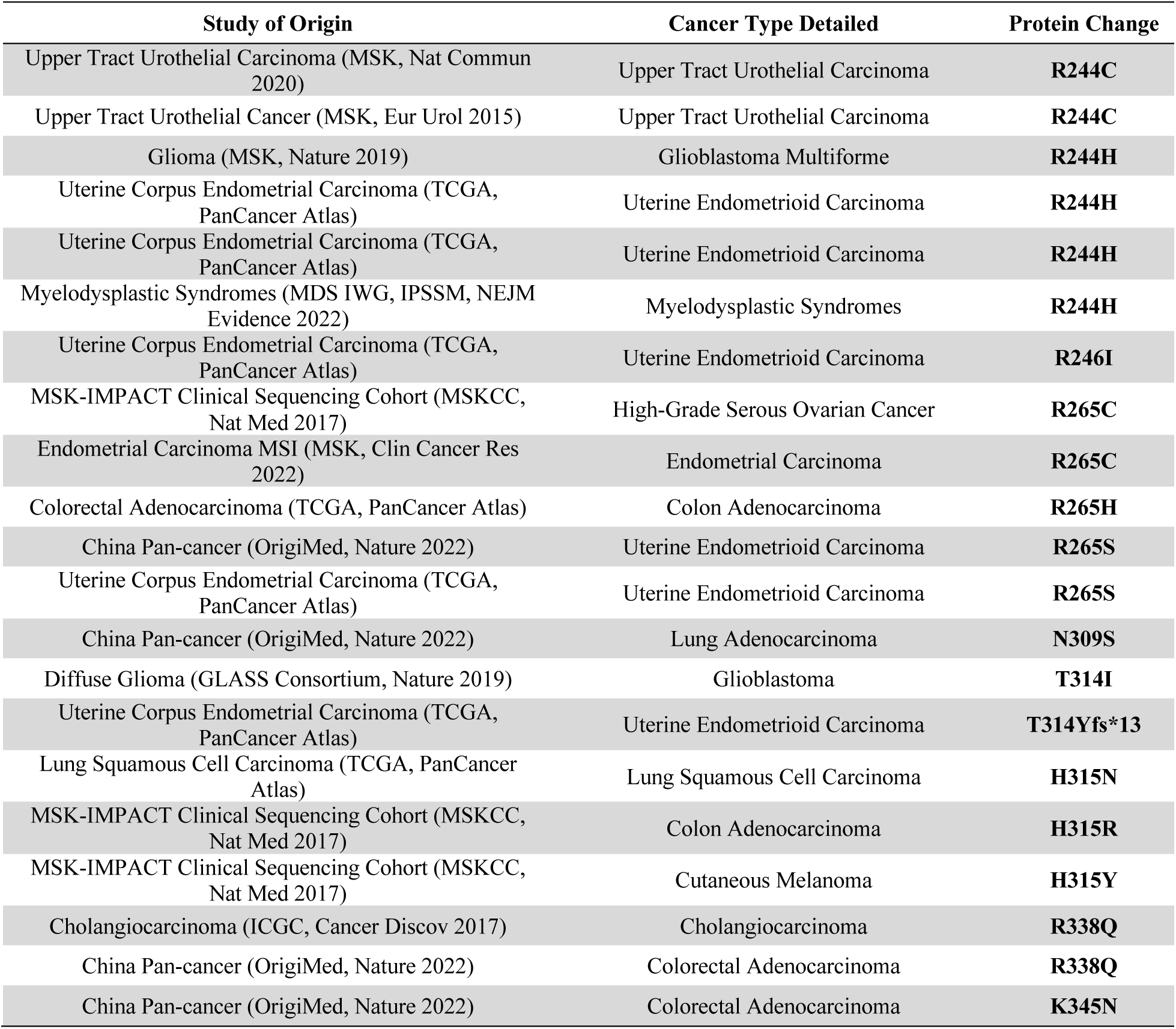
Tabulated cBioPortal mutations to ASXL1. Table showing mutations from cBioPoral to ASXL1 that interface with DNA dyad, DNA exit, and ubiquitin patch. Columns show study of origin, cancer type, and protein change.

**Supplementary Table S4.** Tabulated cBioPortal mutations to BAP1. Table showing mutations from cBioPoral to BAP1 that interface with acidic patch, DNA dyad, DNA exit, and ubiquitin patch. Columns show study of origin, cancer type, and protein change.

| Study of Origin | Cancer Type Detailed | Protein Change |
| --- | --- | --- |
| Adenoid Cystic Carcinoma Project (J Clin Invest 2019) | Adenoid Cystic Carcinoma | **D11Afs\*58** |
| China Pan-cancer (Origimed, Nature 2022) | Intrahepatic Cholangiocarcinoma | **D11N** |
| China Pan-cancer (Origimed, Nature 2022) | Gastric Adenocarcinoma | **D11N** |
| Cholangiocarcinoma (ICGC, Cancer Discov 2017) | Cholangiocarcinoma | **E31del** |
| Ovarian Serous Cystadenocarcinoma (TCGA, PanCancer Atlas) | Serous Ovarian Cancer | **E31del** |
| MSK-IMPACT Clinical Sequencing Cohort (MSKCC, Nat Med 2017) | Cutaneous Melanoma | **E31del** |
| MSK-IMPACT Clinical Sequencing Cohort (MSKCC, Nat Med 2017) | Unclassified Renal Cell Carcinoma | **E31del** |
| China Pan-cancer (Origimed, Nature 2022) | Liver Hepatocellular Carcinoma | **E31del** |
| Adenoid Cystic Carcinoma Project (J Clin Invest 2019) | Adenoid Cystic Carcinoma | **E31\*** |
| Kidney Renal Clear Cell Carcinoma (TCGA, PanCancer Atlas) | Renal Clear Cell Carcinoma | **E31A** |
| Uterine Corpus Endometrial Carcinoma (TCGA, PanCancer Atlas) | Uterine Endometrioid Carcinoma | **E31D** |
| Hepatocellular Carcinomas (INSERM, Nat Genet 2015) | Hepatocellular Carcinoma | **E31Gfs\*41** |
| Kidney Renal Papillary Cell Carcinoma (TCGA, PanCancer Atlas) | Papillary Renal Cell Carcinoma | **E31G** |
| China Pan-cancer (Origimed, Nature 2022) | Renal Clear Cell Carcinoma | **E31G** |
| Bladder Cancer (MSKCC, Eur Urol 2014) | Bladder Urothelial Carcinoma | **E31K** |
| China Pan-cancer (Origimed, Nature 2022) | Perihilar Cholangiocarcinoma | **E31K** |
| Urothelial Carcinoma (BCAN/HCRN 2022) | Bladder Urothelial Carcinoma | **E31Q** |
| Renal Clear Cell Carcinoma (UTokyo, Nat Genet 2013) | Renal Clear Cell Carcinoma with Sarcomatoid Features | **E31V** |
| China Pan-cancer (Origimed, Nature 2022) | Renal Clear Cell Carcinoma | **E31V** |
| MSK-IMPACT Clinical Sequencing Cohort (MSKCC, Nat Med 2017) | Oligodendroglioma | **Y33del** |
| Uveal Melanoma (TCGA, PanCancer Atlas) | Uveal Melanoma | **Y33\*** |
| Combined Hepatocellular and Intrahepatic Cholangiocarcinoma (Peking University, Cancer Cell 2019) | Hepatocellular Carcinoma plus Intrahepatic Cholangiocarcinoma | **Y33\*** |
| Combined Hepatocellular and Intrahepatic Cholangiocarcinoma (Peking University, Cancer Cell 2019) | Hepatocellular Carcinoma plus Intrahepatic Cholangiocarcinoma | **Y33\*** |
| Breast Cancer (METABRIC, Nature 2012 & Nat Commun 2016) | Breast Invasive Ductal Carcinoma | **Y33\*** |
| Stomach Adenocarcinoma (TCGA, PanCancer Atlas) | Diffuse Type Stomach Adenocarcinoma | **Y33\*** |
| Kidney Renal Papillary Cell Carcinoma (TCGA, PanCancer Atlas) | Papillary Renal Cell Carcinoma | **Y33C** |
| China Pan-cancer (Origimed, Nature 2022) | Intrahepatic Cholangiocarcinoma | **Y33Rfs\*36** |
| Cutaneous Squamous Cell Carcinoma (DFCI, Clin Cancer Res 2015) | Cutaneous Squamous Cell Carcinoma | **L35F** |
| Skin Cutaneous Melanoma (TCGA, PanCancer Atlas) | Cutaneous Melanoma | **L35R** |
| Stomach Adenocarcinoma (U Tokyo, Nat Genet 2014) | Stomach Adenocarcinoma | **R56C** |
| MSK-IMPACT Clinical Sequencing Cohort (MSKCC, Nat Med 2017) | Bladder Urothelial Carcinoma | **R56C** |
| Breast Cancer (METABRIC, Nature 2012 & Nat Commun 2016) | Breast Invasive Ductal Carcinoma | **R56C** |
| Breast Cancer (METABRIC, Nature 2012 & Nat Commun 2016) | Breast Invasive Ductal Carcinoma | **R56C** |
| Uterine Corpus Endometrial Carcinoma (TCGA, PanCancer Atlas) | Uterine Endometrioid Carcinoma | **R56C** |
| China Pan-cancer (Origimed, Nature 2022) | Esophageal Squamous Cell Carcinoma | **R56H** |
| Upper Tract Urothelial Carcinoma (MSK, Nat Commun 2020) | Upper Tract Urothelial Carcinoma | **R57Q** |
| MSK-IMPACT Clinical Sequencing Cohort (MSKCC, Nat Med 2017) | Upper Tract Urothelial Carcinoma | **R57Q** |
| Pediatric Brain Cancer (CPTAC/CHOP, Cell 2020) | Medulloblastoma | **R57W** |
| Pediatric Brain Cancer (CPTAC/CHOP, Cell 2020) | Medulloblastoma | **R57W** |
| Skin Cutaneous Melanoma (Yale, Nat Genet 2012) | Cutaneous Melanoma | **R57_R60del** |
| MSK-IMPACT Clinical Sequencing Cohort (MSKCC, Nat Med 2017) | Upper Tract Urothelial Carcinoma | **R59Q** |
| MSK-IMPACT Clinical Sequencing Cohort (MSKCC, Nat Med 2017) | Colorectal Adenocarcinoma | **R59Q** |
| China Pan-cancer (Origimed, Nature 2022) | Gastric Adenocarcinoma | **R59Q** |
| Prostate Adenocarcinoma (SMMU, Eur Urol 2017) | Prostate Adenocarcinoma | **R59Q** |
| Skin Cutaneous Melanoma (TCGA, PanCancer Atlas) | Cutaneous Melanoma | **R59W** |
| Cholangiocarcinoma (ICGC, Cancer Discov 2017) | Cholangiocarcinoma | **R59W** |
| MSK-IMPACT Clinical Sequencing Cohort (MSKCC, Nat Med 2017) | Uveal Melanoma | **R60\*** |
| China Pan-cancer (Origimed, Nature 2022) | Pancreatic Adenocarcinoma | **R60\*** |
| Endometrial Carcinoma MSI (MSK, Clin Cancer Res 2022) | Endometrial Carcinoma | **R60Q** |
| MSK-IMPACT Clinical Sequencing Cohort (MSKCC, Nat Med 2017) | Intrahepatic Cholangiocarcinoma | **C91F** |
| MSK-IMPACT Clinical Sequencing Cohort (MSKCC, Nat Med 2017) | Renal Clear Cell Carcinoma | **C91F** |
| China Pan-cancer (Origimed, Nature 2022) | Renal Clear Cell Carcinoma | **C91F** |
| Mesothelioma (TCGA, PanCancer Atlas) | Pleural Mesothelioma, Epithelioid Type | **C91G** |
| Renal Clear Cell Carcinoma (UTokyo, Nat Genet 2013) | Renal Clear Cell Carcinoma with Sarcomatoid Features | **C91G** |
| MSK-IMPACT Clinical Sequencing Cohort (MSKCC, Nat Med 2017) | Bladder Urothelial Carcinoma | **C91R** |
| China Pan-cancer (Origimed, Nature 2022) | Intrahepatic Cholangiocarcinoma | **C91R** |
| Urothelial Carcinoma (BCAN/HCRN 2022) | Bladder Urothelial Carcinoma | **C91R** |
| Pan-cancer Analysis of Advanced and Metastatic Tumors (BCGSC, Nature Cancer 2020) | Basal Cell Carcinoma | **C91S** |
| China Pan-cancer (Origimed, Nature 2022) | Perihilar Cholangiocarcinoma | **C91W** |
| Pleural Mesothelioma (NYU, Cancer Res 2015) | Pleural Mesothelioma | **C91Y** |
| Urothelial Carcinoma (BCAN/HCRN 2022) | Bladder Urothelial Carcinoma | **C91Y** |
| MSK-IMPACT Clinical Sequencing Cohort (MSKCC, Nat Med 2017) | Cholangiocarcinoma | **R146K** |
| SUMMIT - Neratinib Basket Study (Multi-Institute, Nature 2018) | Cholangiocarcinoma | **R146K** |
| Metastatic Biliary Tract Cancers (SUMMIT - Neratinib Basket Trial, 2022) | Cholangiocarcinoma | **R146K** |
| China Pan-cancer (Origimed, Nature 2022) | Intrahepatic Cholangiocarcinoma | **R146M** |
| Diffuse Glioma (GLASS Consortium, Nature 2019) | Glioblastoma | **X146_splice** |
| Cholangiocarcinoma (ICGC, Cancer Discov 2017) | Cholangiocarcinoma | **X146_splice** |
| Cholangiocarcinoma (ICGC, Cancer Discov 2017) | Cholangiocarcinoma | **X146_splice** |
| Pan-cancer Analysis of Advanced and Metastatic Tumors (BCGSC, Nature Cancer 2020) | Pancreatic Adenocarcinoma | **X146_splice** |
| MSK-IMPACT Clinical Sequencing Cohort (MSKCC, Nat Med 2017) | Intrahepatic Cholangiocarcinoma | **X146_splice** |
| MSK-IMPACT Clinical Sequencing Cohort (MSKCC, Nat Med 2017) | Pleural Mesothelioma | **X146_splice** |
| MSK-IMPACT Clinical Sequencing Cohort (MSKCC, Nat Med 2017) | Uveal Melanoma | **X146_splice** |
| Pan-cancer analysis of whole genomes (ICGC/TCGA, Nature 2020) | Hepatocellular Carcinoma | **X146_splice** |
| China Pan-cancer (Origimed, Nature 2022) | Colorectal Adenocarcinoma | **X146_splice** |
| China Pan-cancer (Origimed, Nature 2022) | Gastric Adenocarcinoma | **X146_splice** |
| China Pan-cancer (Origimed, Nature 2022) | Colorectal Adenocarcinoma | **X146_splice** |
| Liver Hepatocellular Carcinoma (TCGA, PanCancer Atlas) | Hepatocellular Carcinoma | **X146_splice** |
| Clear Cell Renal Cell Carcinoma (DFCI, Science 2019) | Renal Clear Cell Carcinoma | **R227Afs\*4** |
| MSK-IMPACT Clinical Sequencing Cohort (MSKCC, Nat Med 2017) | Prostate Adenocarcinoma | **R227C** |
| MSK-IMPACT Clinical Sequencing Cohort (MSKCC, Nat Med 2017) | Prostate Adenocarcinoma | **R227C** |
| Adenoid Cystic Carcinoma Project (J Clin Invest 2019) | Adenoid Cystic Carcinoma | **R227Hfs\*18** |
| China Pan-cancer (Origimed, Nature 2022) | Gastric Adenocarcinoma | **R227Hfs\*17** |
| Pan-cancer analysis of whole genomes (ICGC/TCGA, Nature 2020) | Lung Squamous Cell Carcinoma | **R227H** |
| China Pan-cancer (Origimed, Nature 2022) | Perihilar Cholangiocarcinoma | **R227H** |
| Upper Tract Urothelial Carcinoma (MSK, Nat Commun 2020) | Upper Tract Urothelial Carcinoma | **R227L** |
| Upper Tract Urothelial Cancer (MSK, Eur Urol 2015) | Upper Tract Urothelial Carcinoma | **R227L** |
| Upper Tract Urothelial Carcinoma (MSK, Nat Commun 2020) | Upper Tract Urothelial Carcinoma | **R227P** |
| Upper Tract Urothelial Cancer (MSK, Eur Urol 2015) | Upper Tract Urothelial Carcinoma | **R227P** |
| Adenoid Cystic Carcinoma Project (J Clin Invest 2019) | Adenoid Cystic Carcinoma | **R227P** |
| Adenoid Cystic Carcinoma Project (J Clin Invest 2019) | Adenoid Cystic Carcinoma | **R227P** |
| MSK-IMPACT Clinical Sequencing Cohort (MSKCC, Nat Med 2017) | Upper Tract Urothelial Carcinoma | **R227P** |
| MSK-IMPACT Clinical Sequencing Cohort (MSKCC, Nat Med 2017) | Bladder Urothelial Carcinoma | **R227P** |
| China Pan-cancer (Origimed, Nature 2022) | Thymic Carcinoma | **R227P** |
| China Pan-cancer (Origimed, Nature 2022) | Pancreatic Adenocarcinoma | **R227P** |
| Urothelial Carcinoma (BCAN/HCRN 2022) | Bladder Urothelial Carcinoma | **R227P** |
| China Pan-cancer (Origimed, Nature 2022) | Head and Neck Squamous Cell Carcinoma | **L230P** |
| Renal Clear Cell Carcinoma (UTokyo, Nat Genet 2013) | Renal Clear Cell Carcinoma with Sarcomatoid Features | **L230Q** |
| Uterine Corpus Endometrial Carcinoma (TCGA, PanCancer Atlas) | Uterine Endometrioid Carcinoma | **D672N** |
| Breast Cancer (METABRIC, Nature 2012 & Nat Commun 2016) | Breast Invasive Ductal Carcinoma | **R699P** |
| Uveal Melanoma (QIMR, Oncotarget 2016) | Uveal Melanoma | **R699Qfs\*6** |
| China Pan-cancer (Origimed, Nature 2022) | Pancreatic Neuroendocrine Tumor | **R699Q** |
| MSK-IMPACT Clinical Sequencing Cohort (MSKCC, Nat Med 2017) | Colorectal Adenocarcinoma | **R699W** |
| Breast Cancer (METABRIC, Nature 2012 & Nat Commun 2016) | Breast Invasive Ductal Carcinoma | **R699W** |
| China Pan-cancer (Origimed, Nature 2022) | Lung Squamous Cell Carcinoma | **R700L** |
| Pan-cancer analysis of whole genomes (ICGC/TCGA, Nature 2020) | Head and Neck Squamous Cell Carcinoma | **R700Q** |
| China Pan-cancer (Origimed, Nature 2022) | Small Bowel Adenocarcinoma | **R700Q** |
| Uterine Corpus Endometrial Carcinoma (TCGA, PanCancer Atlas) | Uterine Endometrioid Carcinoma | **R700Q** |
| Stomach Adenocarcinoma (TCGA, PanCancer Atlas) | Stomach Adenocarcinoma | **R700Q** |
| Colon Cancer (CPTAC-2 Prospective, Cell 2019) | Colon Adenocarcinoma | **R700W** |
| China Pan-cancer (Origimed, Nature 2022) | Colorectal Adenocarcinoma | **R700W** |
| Uterine Corpus Endometrial Carcinoma (TCGA, PanCancer Atlas) | Uterine Endometrioid Carcinoma | **R700W** |
| Uterine Corpus Endometrial Carcinoma (TCGA, PanCancer Atlas) | Uterine Endometrioid Carcinoma | **R700W** |
| Non-Small Cell Lung Cancer (TRACERx, NEJM & Nature 2017) | Non-Small Cell Lung Cancer | **R701C** |
| Non-Small Cell Lung Cancer (TRACERx, NEJM & Nature 2017) | Non-Small Cell Lung Cancer | **R701C** |
| Non-Small Cell Lung Cancer (TRACERx, NEJM & Nature 2017) | Non-Small Cell Lung Cancer | **R701C** |
| MSK-IMPACT Clinical Sequencing Cohort (MSKCC, Nat Med 2017) | Adrenocortical Carcinoma | **R701C** |

## References and Notes

1. A. P. Szczepanski, L. Wang, Emerging multifaceted roles of BAP1 complexes in biological processes. Cell Death Discovery 7, (2021).

2. J. C. Scheuermann et al., Histone H2A deubiquitinase activity of the Polycomb repressive complex PR-DUB. Nature 465, 243–247 (2010).

3. D. D. Sahtoe, W. J. Van Dijk, R. Ekkebus, H. Ovaa, T. K. Sixma, BAP1/ASXL1 recruitment and activation for H2A deubiquitination. Nature Communications 7, 10292 (2016).

4. L. Sanchez-Pulido, L. Kong, C. P. Ponting, A common ancestry for BAP1 and Uch37 regulators. Bioinformatics 28, 1953–1956 (2012).

5. N. A. Fursova et al., BAP1 constrains pervasive H2AK119ub1 to control the transcriptional potential of the genome. Genes & Development 35, 749–770 (2021).

6. E. Conway et al., BAP1 enhances Polycomb repression by counteracting widespread H2AK119ub1 deposition and chromatin condensation. Molecular Cell 81, 3526–3541.e3528 (2021).

7. J. N. Kuznetsov et al., BAP1 regulates epigenetic switch from pluripotency to differentiation in developmental lineages giving rise to BAP1-mutant cancers. Science Advances 5, eaax1738 (2019).

8. A. Campagne et al., BAP1 complex promotes transcription by opposing PRC1-mediated H2A ubiquitylation. Nature Communications 10, (2019).

9. Y. B. Schwartz, V. Pirrotta, Polycomb silencing mechanisms and the management of genomic programmes. Nature Reviews Genetics 8, 9–22 (2007).

10. L.-H. Wang, M. A. E. Aberin, S. Wu, S.-P. Wang, The MLL3/4 H3K4 methyltransferase complex in establishing an active enhancer landscape. Biochemical Society Transactions 49, 1041–1054 (2021).

11. L. Wang et al., Resetting the epigenetic balance of Polycomb and COMPASS function at enhancers for cancer therapy. Nature Medicine 24, 758–769 (2018).

12. I. H. Ismail et al., Germline Mutations in BAP1 Impair Its Function in DNA Double-Strand Break Repair. Cancer Research 74, 4282–4294 (2014).

13. O. Abdel-Wahab et al., Deletion of Asxl1 results in myelodysplasia and severe developmental defects in vivo. Journal of Experimental Medicine 210, 2641–2659 (2013).

14. M. Cigognetti et al., BAP1 (BRCA1-associated protein 1) is a highly specific marker for differentiating mesothelioma from reactive mesothelial proliferations. Modern Pathology 28, 1043–1057 (2015).

15. M. Cheung, J. R. Testa, BAP1, a tumor suppressor gene driving malignant mesothelioma. Translational Lung Cancer Research 6, 270–278 (2017).

16. F. Matheus et al., Pathological ASXL1 Mutations and Protein Variants Impair Neural Crest Development. Stem Cell Reports 12, 861–868 (2019).

17. L. Shahriyari, M. Abdel-Rahman, C. Cebulla, BAP1 expression is prognostic in breast and uveal melanoma but not colon cancer and is highly positively correlated with RBM15B and USP19. PLoS One 14, e0211507 (2019).

18. G. Stålhammar, T. R. O. See, S. Phillips, S. Seregard, H. E. Grossniklaus, Digital Image Analysis of BAP-1 Accurately Predicts Uveal Melanoma Metastasis. Translational Vision Science & Technology 8, 11 (2019).

19. I. De et al., Structural Basis for the Activation of the Deubiquitinase Calypso by the Polycomb Protein ASX. Structure 27, 528–536.e524 (2019).

20. M. Foglizzo et al., A bidentate Polycomb Repressive-Deubiquitinase complex is required for efficient activity on nucleosomes. Nature Communications 9, (2018).

21. D. Sahtoe, Danny et al., Mechanism of UCH-L5 Activation and Inhibition by DEUBAD Domains in RPN13 and INO80G. Molecular Cell 57, 887–900 (2015).

22. L. Long, M. Furgason, T. Yao, Generation of nonhydrolyzable ubiquitin-histone mimics. Methods 70, 134–138 (2014).

23. H. Stark, GraFix: stabilization of fragile macromolecular complexes for single particle cryo-EM. Methods Enzymol 481, 109–126 (2010).

24. R. Evans, et al., Protein complex prediction with AlphaFold-Multimer. bioRxiv, 2021.2010.2004.463034 (2022).

25. D. Komander, M. Rape, The ubiquitin code. Annu Rev Biochem 81, 203–229 (2012).

26. M. I. Valencia-Sánchez et al., Regulation of the Dot1 histone H3K79 methyltransferase by histone H4K16 acetylation. Science 371, eabc6663 (2021).

27. E. J. Worden, X. Zhang, C. Wolberger, Structural basis for COMPASS recognition of an H2B-ubiquitinated nucleosome. eLife 9, (2020).

28. H. Peng et al., Familial and Somatic BAP1 Mutations Inactivate ASXL1/2-Mediated Allosteric Regulation of BAP1 Deubiquitinase by Targeting Multiple Independent Domains. Cancer Res 78, 1200–1213 (2018).

29. C. A. Davey, D. F. Sargent, K. Luger, A. W. Maeder, T. J. Richmond, Solvent mediated interactions in the structure of the nucleosome core particle at 1.9 a resolution. J Mol Biol 319, 1097–1113 (2002).

30. V. Kasinath et al., JARID2 and AEBP2 regulate PRC2 in the presence of H2AK119ub1 and other histone modifications. Science 371, (2021).

31. M. I. Valencia-Sanchez et al., Structural Basis of Dot1L Stimulation by Histone H2B Lysine 120 Ubiquitination. Mol Cell 74, 1010–1019 e1016 (2019).

32. R. K. McGinty, S. Tan, Recognition of the nucleosome by chromatin factors and enzymes. Curr Opin Struct Biol 37, 54–61 (2016).

33. R. K. Mcginty, R. C. Henrici, S. Tan, Crystal structure of the PRC1 ubiquitylation module bound to the nucleosome. Nature 514, 591–596 (2014).

34. R. D. Makde, J. R. England, H. P. Yennawar, S. Tan, Structure of RCC1 chromatin factor bound to the nucleosome core particle. Nature 467, 562–566 (2010).

35. A. J. Barbera et al., The Nucleosomal Surface as a Docking Station for Kaposi’s Sarcoma Herpesvirus LANA. Science 311, 856–861 (2006).

36. K. J. Armache, J. D. Garlick, D. Canzio, G. J. Narlikar, R. E. Kingston, Structural basis of silencing: Sir3 BAH domain in complex with a nucleosome at 3.0 A resolution. Science 334, 977–982 (2011).

37. E. J. Worden, N. A. Hoffmann, C. W. Hicks, C. Wolberger, Mechanism of Cross-talk between H2B Ubiquitination and H3 Methylation by Dot1L. Cell 176, 1490–1501.e1412 (2019).

38. M. Carbone et al., Biological Mechanisms and Clinical Significance of BAP1 Mutations in Human Cancer. Cancer Discovery 10, 1103–1120 (2020).

39. H. Yang et al., Gain of function of ASXL1 truncating protein in the pathogenesis of myeloid malignancies. Blood 131, 328–341 (2018).

40. L. Wang et al., Epigenetic targeted therapy of stabilized BAP1 in ASXL1 gain-of-function mutated leukemia. Nature Cancer 2, 515–526 (2021).

41. A. Skrajna et al., Comprehensive nucleosome interactome screen establishes fundamental principles of nucleosome binding. Nucleic Acids Research 48, 9415–9432 (2020).

42. D. Grau et al., Structures of monomeric and dimeric PRC2:EZH1 reveal flexible modules involved in chromatin compaction. Nature Communications 12, (2021).

43. M. Dombrowski, M. Engeholm, C. Dienemann, S. Dodonova, P. Cramer, Histone H1 binding to nucleosome arrays depends on linker DNA length and trajectory. Nature Structural & Molecular Biology 29, 493–501 (2022).

44. B.-R. Zhou et al., Distinct Structures and Dynamics of Chromatosomes with Different Human Linker Histone Isoforms. Molecular Cell 81, 166–182.e166 (2021).

45. R. Meas, P. Mao, Histone ubiquitylation and its roles in transcription and DNA damage response. DNA Repair 36, 36–42 (2015).

46. P. N. Dyer et al., Reconstitution of nucleosome core particles from recombinant histones and DNA. Methods Enzymol 375, 23–44 (2004).

47. L. F. Schachner et al., Decoding the protein composition of whole nucleosomes with Nuc-MS. Nat Methods 18, 303–308 (2021).

48. M. R. Marunde et al., Nucleosome conformation dictates the histone code. bioRxiv, (2022).

49. X. Bi, R. Yang, X. Feng, D. Rhodes, C. F. Liu, Semisynthetic UbH2A reveals different activities of deubiquitinases and inhibitory effects of H2A K119 ubiquitination on H3K36 methylation in mononucleosomes. Org Biomol Chem 14, 835–839 (2016).

50. Y. S. Choi et al., High-affinity free ubiquitin sensors for quantifying ubiquitin homeostasis and deubiquitination. Nat Methods 16, 771–777 (2019).

51. X. Li et al., Electron counting and beam-induced motion correction enable near-atomic-resolution single-particle cryo-EM. Nat Methods 10, 584–590 (2013).

52. A. Cheng et al., Leginon: New features and applications. Protein Sci 30, 136–150 (2021).

53. S. Q. Zheng et al., MotionCor2: anisotropic correction of beam-induced motion for improved cryo-electron microscopy. Nat Methods 14, 331–332 (2017).

54. J. Zivanov et al., New tools for automated high-resolution cryo-EM structure determination in RELION-3. Elife 7, (2018).

55. A. Punjani, J. L. Rubinstein, D. J. Fleet, M. A. Brubaker, cryoSPARC: algorithms for rapid unsupervised cryo-EM structure determination. Nat Methods 14, 290–296 (2017).

56. K. Zhang, Gctf: Real-time CTF determination and correction. J Struct Biol 193, 1–12 (2016).

57. S. Vijay-Kumar, C. E. Bugg, W. J. Cook, Structure of ubiquitin refined at 1.8 A resolution. J Mol Biol 194, 531–544 (1987).

58. E. F. Pettersen et al., UCSF Chimera--a visualization system for exploratory research and analysis. J Comput Chem 25, 1605–1612 (2004).

59. P. D. Adams et al., PHENIX: a comprehensive Python-based system for macromolecular structure solution. Acta Crystallogr D Biol Crystallogr 66, 213–221 (2010).

60. P. Emsley, K. Cowtan, Coot: model-building tools for molecular graphics. Acta Crystallogr D Biol Crystallogr 60, 2126–2132 (2004).

61. E. F. Pettersen et al., UCSF ChimeraX: Structure visualization for researchers, educators, and developers. Protein Sci 30, 70–82 (2021).

62. Schrodinger, LLC. (2015).

63. M. D. Wilson et al., The structural basis of modified nucleosome recognition by 53BP1. Nature 536, 100–103 (2016).

